# Asymmetric dimethylarginine positively modulates Calcium-Sensing Receptor signalling to promote lipid accumulation and adiposity

**DOI:** 10.1101/2022.07.26.501411

**Authors:** Laura Dowsett, Lucie Duluc, Erin Higgins, Fatmah Alghamdi, Walter Fast, Ian P. Salt, James Leiper

## Abstract

Irreversible methylation of arginine residues generates asymmetric dimethylarginine (ADMA). ADMA is a competitive inhibitor of nitric oxide (NO) synthase and an independent risk factor for cardiovascular disease. Plasma ADMA concentrations increase with obesity and fall following weight loss. Here, we demonstrate that ADMA drives lipid accumulation through a newly identified NO-independent pathway *via* the amino-acid sensitive calcium-sensing receptor (CaSR). ADMA treatment of 3T3-L1 and HepG2 cells activates mTOR signalling and upregulates a suite of lipogenic genes with an associated increase in triglyceride content. Pharmacological blockade of CaSR inhibits ADMA driven lipid accumulation and ADMA treatment potentiates CaSR signalling via both G_q_ and G_i/o_ pathways. Impairment of ADMA metabolism in adipocytes *in vivo*, by dimethylamine dimethylaminohydrolase-1 (DDAH1) deletion, increases visceral adiposity and adipocyte hypertrophy. This study identifies a signalling mechanism for ADMA as an endogenous ligand of the G protein-coupled receptor CaSR that potentially contributes to the impact of ADMA in cardiometabolic disease.

## Introduction

Methylarginines are formed via the methylation of arginine residues by protein arginine methyltransferases (PRMT) (Bedford and Clarke, 2009). The PRMT family consists of nine enzymes each encoded by a separate gene. These methylate histone proteins as a key step in transcriptional regulation and a wider range of non-histone proteins involved in protein synthesis. All PRMT enzymes are capable of forming L-N^G^ monomethylated arginine (L-NMMA); while type I PRMTs, predominantly PRMT 1, form asymmetric dimethylarginine (ADMA) and type II PRMTs, predominantly PRMT 5, form symmetric dimethylarginine (SDMA). Unlike other post-translational modifications, arginine methylation is irreversible and therefore, proteolysis generates a constant flow of free methylarginines. These are initially released into the cytosol from where they may exit the cell to the circulation and ultimately be cleared by the kidney (Vallance *et al*., 1992a). Previous studies, from our group and others, have demonstrated that asymmetrically methylated methylarginines are competitive inhibitors of all three isoforms of nitric oxide synthase (Vallance *et al*., 1992b; Cardounel and Zweier, 2002; Leiper *et al*., 2007). In contrast, SDMA has no inhibitory action on NOS. As ADMA concentrations are ∼10-fold higher than that of L-NMMA it is thought ADMA is the most significant endogenously produced inhibitor of NO synthesis.

Relatively modest increases in ADMA in both humans and experimental models are associated with profound disruption of cardiovascular homeostasis and have been linked to an increased risk of hypertension, atherosclerosis and stroke (Boger *et al*., 1998; Leiper *et al*., 2007; Lambden *et al*., 2015). Further clinical studies have associated elevated plasma ADMA concentrations with obesity and the metabolic syndrome (Eid *et al*., 2004; Kocak *et al*., 2011; McLaughlin *et al*., 2006; Palomo *et al*., 2011). Concentrations vary widely between studies but on average increase by 0.3 μM from ∼0.95 μM to ∼1.3 μM. ADMA accumulates in the visceral adipose tissue of obese subjects (Assar *et al*., 2016) and circulating ADMA levels decrease upon weight loss although causal mechanisms have not been identified (McLaughlin *et al*., 2006; Krzyzanowska *et al*., 2004). Interestingly however, NO levels are also increased in obesity and correlate with both BMI and adiposity (Choi *et al*., 2001) with higher expression of eNOS and iNOS in the adipose tissue of obese individuals (Elizalde *et al*., 2000). These observations might suggest additional roles of ADMA in adipose tissue that are independent of NOS. Therefore, in this study we set out to test the hypothesis that ADMA regulates adipocyte function and physiology via a NO-independent mechanism. Upon establishing that NO-independent pathways exist in both adipocytes and hepatocytes we then went on to test the hypothesis that ADMA may act via the amino-acid sensitive calcium-sensing receptor.

## Methods

### Contact for reagents and resource sharing

Further information and requests for resources and reagents can be directed to, James Leiper or Laura Dowsett.

### Experimental Models

#### Mice

All studies were conducted in accordance with UK Home Office Animals (Scientific Procedures) Act 1986 and institutional guidelines with the Imperial College ethics review board approval.

The DDAH1 floxed mouse (DDAH1^fl/fl^) was developed for us by GenOway (Lyon, France). Cre-LoxP site were inserted into the DDAH1 gene on either side of exon one (Dowsett *et al*., 2015). The adipocyte specific DDAH1 knockout mouse (DDAH1^Ad-/-^) was developed by crossing the DDAH1^fl/fl^ with the Adiponectin driven cre mouse strain (FVB-Tg(Adipoq-cre)1Evdr/J) developed by Evan Rosen *et al*. (Eguchi *et al*., 2011) DDAH1^fl/fl^ mice were used as controls. All mice were derived on a C57/bl6 background. Mice were maintained in house on a fixed light/dark cycle and fed *ad libitum*. Male DDAH1^Ad-/-^ and littermate controls were weighed every two weeks from six weeks of age. Fat and lean mass was assessed using a body composition analysis Scanner (EchoMRI, Houston, USA). Whole body calorimetry was measured using the CLAMS: Comprehensive lab Animal Monitoring System (Columbus Instruments, Columbus, USA). Mice were acclimatised in the system for 8 hours before data collection for the following 72 hours.

### Cell Lines

#### Immortalized Cell Lines

The murine pre-adipocyte cell line 3T3-L1 was purchased from American Type Culture Collection (ATCC CL-173) and subcultured according to manufacturer’s instruction at 37 °C in a humidified atmosphere with 5% CO_2_. To stimulate differentiation to mature adipocytes, 2 days post-confluent cells were exposed to high glucose (4.5 g/L) Dulbecco’s modified Eagle’s Medium (DMEM) supplemented with 10% fetal bovine serum (FBS), 1x penicillin-streptomycin and 2mM L-glutamine and containing 10 µg/ml insulin (Sigma-Aldrich, I9278), 1 µM dexamethasone (Sigma-Aldrich, D1756) and 0.5 mM iso-butyl-methylxanthine (Sigma-Aldrich, I5879). Induction media was replaced two days later with media containing 10 µg/ml insulin only. Cells were then cultured a further 4 days in high glucose DMEM. Adipocytes were therefore considered mature at 10 days post induction.

The human hepatoma cell line HepG2 was purchased from American type Culture Collection (ATCC HB-8065) Cells were maintained at 37 °C in a humidified atmosphere with 5% CO_2_ in DMEM supplemented with 10% FBS, 2 mM glutamine and 1% penicillin-streptomycin.

#### Primary Murine Adipocytes

Primary adipocytes were isolated as developed by Rodbell (Rodbell, 1964). Epididymal fat pads were removed from 22 week old mice. Fat pads were washed in PBS and dissected to remove larger blood vessels. The remaining tissue was digested in low-glucose (1 mg/L) DMEM supplemented with 100 mM HEPES, 1.5% BSA and 0.2% collagenase II, with gentle shaking at 37°C for 1 hour. Cells were then centrifuged for 2 minutes at 400 x g; the floating fat cells were collected, washed in PBS and processed for future analysis.

## Method Details

### Cell Culture Treatments

#### 3T3-L1 Treatments

Cell size experiments in mature 3T3-L1 adipocytes were performed for 72 hours with either ADMA (1, 3 and 10 µM, Calbiochem, 311203), SDMA (10 µM, Calbiochem, 311204), PB-ITU (20 µM, Sigma-Aldrich), L-NAME (1 mM, Sigma-Aldrich, N5751) treatment. Experiments to assess mRNA and protein levels were performed over 48 hours including those to explore mTOR inhibition by rapamycin (10 nM, Sigma-Aldrich, R8781), and effects by NO donors S-Nitroso-N-acetyl-DL-penicillamine (SNAP, 100 µM Calbiochem, 487910) and 8-Bromo-Guanosine 3’,5’-cyclic Monophosphate (8-bromo-cGMP, 100 µM Calbiochem 203820). Actinomycin D (5 µg/ml, Sigma-Aldrich, A9415) was used to assess RNA transcription over 30 hours. The role of CaSR was assessed by the agonist Cinacalcet hydrochloride (50 nM, 6170/10) and inhibitors NPS-2143 hydrochloride (10 µM, Tocris, 3626/10), Calhex 231 hydrochloride (1 µM, Tocris, 4387/10) over 48 hours.

#### HepG2 Treatments

HepG2 cells treatments were performed as above with the 3T3-L1 cells but over an 18 hour time course for studies exploring mRNA expression.

### Metabolic Assays

#### Triglycerides and cholesterol quantification

Triglyceride content was measured using Triglycerides Quantification Assay kit from Abcam (ab65336) and total cholesterol content was assessed with Amplex Red Cholesterol Assay Kit from Invitrogen (A12216) according to manufacturer’s instructions. Free fatty acids were measured using an enzyme-based method from Abcam (ab65341).

#### SREBP -1 Transcription Factor Assay

Fully differentiated 3T3-L1 cells were treated for 6 hours with 10µM ADMA. Nuclear extracts were made using the nuclear extraction kit from Abcam (ab113474). Nuclear SREBP-1 was detected using the SREBP-1 transcription factor assay kit from Abcam (ab133125) which uses an ELISA based approach to detect the change in nuclear SREBP-1 in a 96-well format.

#### Insulin Sensitivity

Glucose uptake was measured by the uptake of 2-[^3^H]-deoxy-D-glucose as described previously (Boyle *et al*., 2011). Briefly, 3T3-L1 adipocytes were stimulated with 10 μM ADMA for 48 hours, the medium removed and replaced with Krebs-Ringer-phosphate (KRP (128 mM NaCl, 4.7 mM KCl, 5 mM NaH_2_PO_4_ (pH 7.4), 1.2 mM MgSO_4_, 2.5 mM CaCl_2_, 1% (w/v) BSA, 3 mM glucose)) buffer containing the same concentration of ADMA. After incubation for 1 h at 37°C, cells were incubated in the presence or absence of 0-100 nM insulin for 12 min and transport was initiated by adding [^3^H]-2-deoxyglucose (50 μM, 2 μCi/ml). Cells were incubated for 3 min, after which cells were rapidly washed in ice-cold PBS, air-dried, and then solubilized in 1% (v/v) Triton X-100. The radioactivity associated with the cells was determined by liquid scintillation spectrophotometry. Non-specific cell-associated radioactivity was determined in parallel incubations performed in the presence of 10 μM cytochalasin B.

### Molecular Biology Assays

#### RNA isolation and Rt-qPCR analysis

RNA was extracted and isolated from both cell cultures and mouse tissues using Qiagen’s Universal RNeasy Kit. Mouse tissues were homogenised using a tissue-lyser (Qiagen). cDNA was synthesised from total RNA with iScript cDNA Synthesis kit (Bio-rad). Real-time PCR’s were run on the 7900 Fast Real-time PCR system (Applied Biosciences) using SYBR green Supermix (Bio-rad). Fold changes in mRNA expression were calculated from at least three independent experiments using the standard curve method and normalised to 18S gene expression. Primer sequences are provided in Supplementary Table 1.

#### Western blotting

Following protein electrophoresis and protein transfer, the membranes were probed with primary antibodies from the following sources: mTOR antibody (Cell Signalling, #2972), total AKT (Cell Signalling, #9272), phospho-AKT, Ser^473^ (Cell Signalling, #4060), goat monoclonal DDAH1 and DDAH2 antibodies both developed in house^24^, CaSR (Abcam, ab137408)). Loading controls used were pan-actin (Cell Signalling #8456), and vinculin (Invitrogen #700062). Primary antibodies were used at dilution 1:1000 and secondary antibodies at dilution 1:5000. The relative intensity of the immune-reactive bands was determined by densitometry using Li-COR Image Studio Software. The results were normalised to the loading protein and expressed as fold change over control. Untreated control samples were included in every experiment and all experiments were repeated at least 3 times.

#### Immunohistochemistry

Cells were plated on coverslips, cultured and then fixed with 4% formaldehyde solution in PBS for 10 minutes at room temperature, washed in PBS for 5 minutes and incubated in 3% BSA, 0.1% Triton-X100 blocking buffer in PBS for 1 hour at room temperature. Rabbit anti-CaSR antibody (Abcam #137408) was incubate 1:200 overnight in 3% BSA at 4°C before washing with 0.1% Triton -X100 in PBS. Secondary antibodies (Alexafluor 488, ThermoFisher) were incubated for 2 hours at room temperature before washing and mounting with Prolong Diamond Antifade with DAPI (ThemoFisher).

#### Bodipy

Cells were cultured on coverslips, stimulated with various treatments and then fixed with 4% formaldehyde solution in PBS for 10 minutes at room temperature, washed in PBS for 5 minutes and incubated in PBS containing 1 ug/ml Bodipy (ThermoFisher, D3921) for 15 minutes upon agitation and in the dark. Coverslips were washed three times in PBS and mounted in Prolong Diamond with DAPI mounting media. Pictures were taken under a confocal laser scanning fluorescence microscope (Leica TCS SP5). Cell size was determined using the NIH ImageJ Software package. Three images were captured per a coverslip. Starting in the bottom right for each image 30 cells (or all cells if less than 30) were analysed. Average lipid area was assessed by thresholding of total BODIPY area and divided by the number of lipid laden cells. Data was plotted as an average of 3 images per experiment.

#### Epididymal fat pad staining

Epididymal fat pads were fixed in 4% PFA overnight and then stored in 50% ethanol until embedding in paraffin and sectioning. To assess adipocyte size the tissue was stained with Elastin Van Geisen Stain and all complete cells within each image was measured. Macrophage infiltration was measured by counting the number of Galectin-3 positive cells (Cedarlane). Analysis was performed using Image J software.

### Calcium-Sensing Receptor Assays

#### Flp-In T-REX Hek293-CaSR Cells

Inducible CaSR overexpressing cells were generated using the Flp-In T-REx 293 cell line using the protocol developed by Ward et. al^19^. The N-terminal Myc-tagged human CaSR was inserted by Genescript into the pcDNA5-FRT-TO plasmid. The pcDNA5-FRT-TO and the pOG44 plasmids were transfected using Lipofectamine 2000 (ThermoFisher). Positive cells were selected using hygromycin B. CaSR expression was induced by 0.5 µg/ml doxycycline treatment for 24 hours.

#### Intracellular Ca^2+^ Imaging

Hek293-CaSR cells were incubated at 37°C with 1 µM Cal520 (Abcam, ab171868) in Optimem Reduced Serum I media (HEPES, 2.4 g/l Sodium Bicarbonate, L-glutamine) containing relevant treatments for 1 hour before imaging. Cells were then washed and then incubated in Ca^2+^ - free HEPES buffer (130 mM NaCl, 5 mM KCl, 1 mM MgCl_2_, 20 mM HEPES, 10 mM Glucose, pH7.4) containing relevant treatments. Gd^3+^ was used as a CaSR agonist (0.01 mM-5 mM), NPS-2143 was used as a CaSR antagonist (10 µM). Changes in cytosolic Ca^2+^ were imaged using a Zeiss ZenPro inverted microscope; and FITC fluorescence determined using the physiology definition programme as part of the Zen Software (Zeiss Microscopy GmbH, Germany). Data are represented as the change in fluorescence (FΔ) over the fluorescence at baseline (F0) normalised to the maximum control response to each experiment.

#### Cyclic AMP Analysis

HEK293-CaSR were plated into 96 well plates and treated with doxycycline 24 hours prior to the experiment to induce CaSR expression. Cells were treated with 0.5 mM IBMX for 30 minutes to block phosphodiesterase activity. Cells were then incubated in Optimem reduced serum media with relevant treatments for 15 minutes; 1 µM forskolin was used to stimulate cAMP production. cAMP concentration was assessed using an ELISA kit (Cell Signalling, #4339) as per manufacturers protocol. Data are presented normalised to the maximal forskolin induced response.

#### Quantification and Statistical analysis

Statistical analysis was performed using GraphPad Prism 6 Software. Comparisons were carried out with unpaired two-tailed Student’s *t-*test, One-way and Two-way ANOVA with Bonferroni post-hoc tests as appropriate. Dose response curves were assessed by non-linear regression. Statistical significance was accepted for *P*<0.05. Data are expressed as ± SEM. All *in vitro* experiments are a mean of at least three independent experiments, numbers for each experiment are confirmed in the figure legends.

**Table 1:**
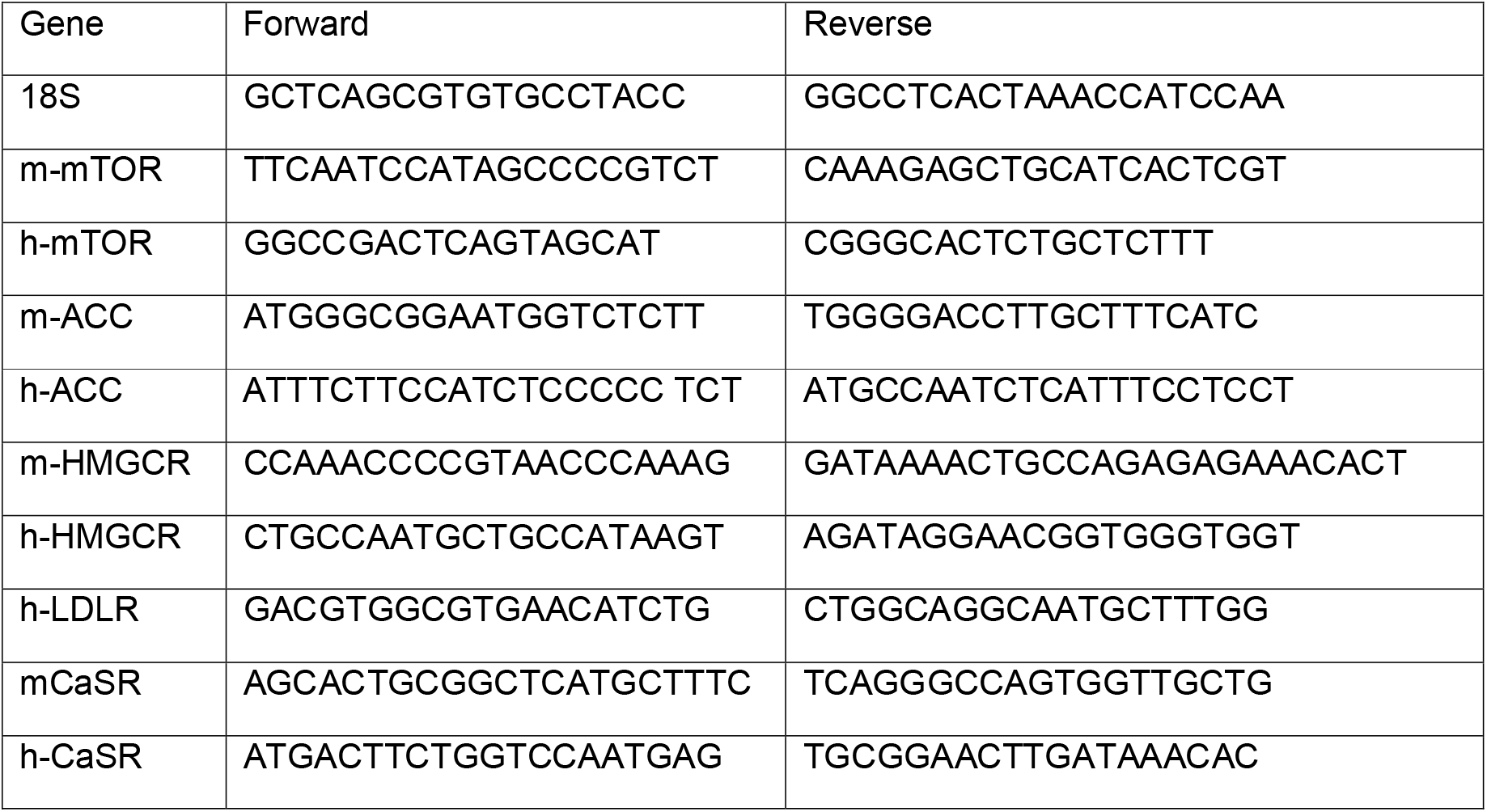
qPCR primer sequences

## Results

### ADMA induces lipid accumulation in 3T3-L1 cells through a NO-independent pathway

To evaluate the consequences of chronic ADMA exposure on adipocyte function we utilised the mouse 3T3-L1 cell line as a model for mature adipocytes. 3T3-L1 fibroblasts were differentiated to lipid laden cells through the addition of insulin, dexamethasone and isobutyl-methylxanthine. 3T3-L1 cells were considered fully differentiated 10 days post-induction, at which point they were treated with ADMA for an additional 72h. ADMA concentrations were chosen to reflect those occurring in disease. 3T3-L1 cells treated with 1 and 3 µM ADMA showed significant cellular hypertrophy compared to those cultured in control media (Fig. 1a and b); whereas SDMA (10 μM) had no effect. Given the currently understood mechanism of action of ADMA is as a competitive inhibitor of NOS we assessed the effect of two structurally distinct synthetic NOS inhibitors N(g)-nitro-L-arginine methyl ester (L-NAME, 1 mM) and 1,3-PBI-TU, Dihydrobromide (20 μM) at concentrations to maximally block NO production. Interestingly, these did not cause adipocyte hypertrophy suggesting that the effect of ADMA on cell size may be independent of NOS inhibition. Taken together these data suggest that a NO independent mechanism specific for ADMA drives adipocytes hypertrophy in 3T3-L1 cells.

**Fig. 1:**
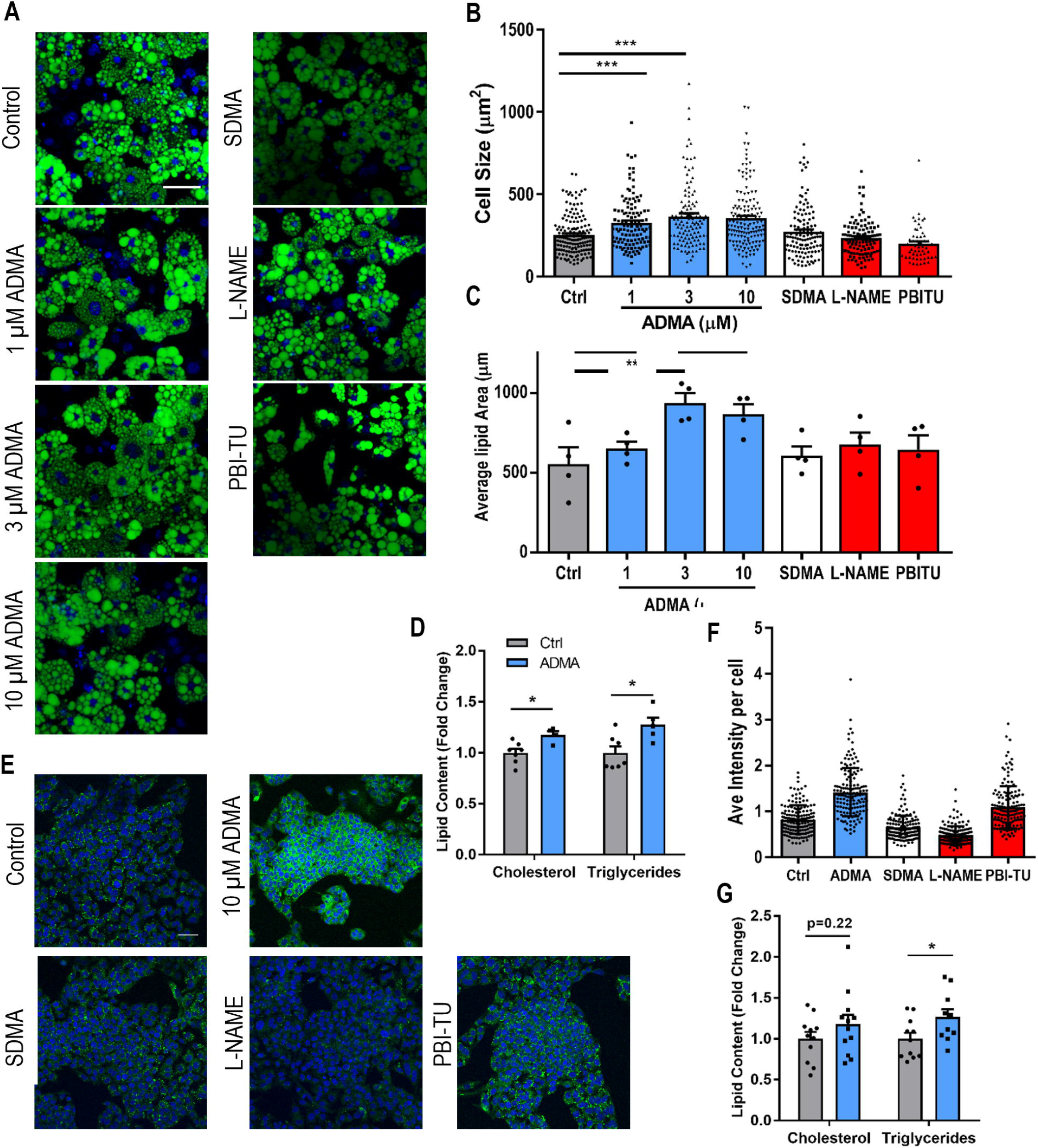
ADMA promotes lipid accumulation through a NO-independent mechanism in 3T3-L1 and HepG2 cells. **(a - c)** 3T3-L1 cells were treated for 72 hours in DMEM supplemented with ADMA, 10 µM SDMA, 1 mM L-NAME and 20 µM PBI-TU were **(a)** stained with BODIPY, (representative images from 3 experiments, scale bar 50 µm) and measured to establish **(b)** cell area and **(c)** the average lipid area per a cell (155 cells from 4 independent experiments). **(d)** 3T3-L1 cholesterol and triglyceride content following 10µM ADMA treatment for 72 hours (N=7). **(e)** HepG2 cells treated for 18 hours with 10µM ADMA, 10µM SDMA, 1mM L-NAME and 20µM PBI-TU and were stained with BODIPY for lipid accumulation; **(f)** lipid accumulation was established through average fluorescent intensity of BODIPY staining per cell. **(g)** HepG2 cholesterol and triglyceride content following 10 µM ADMA treatment for 18 hours (N=11). Analysis by one-way ANOVA followed by multiple comparisons test (Bonferroni) in **b, c** and **f**; and by two-tailed student’s t-test in **d** and **g**. * P<0.05, **P<0.01, ***P<0.001. Data presented are mean ± S.E.M.

In order to identify the potential mechanism through which ADMA is acting we first established whether cellular hypertrophy was due to increased lipid content. Adipocytes treated with 3 µM and 10 µM ADMA have increased lipid area per a cell (Fig. 1c) with a rise in intracellular cholesterol and triglyceride (Fig. 1d). Given that an increase in the proportion of differentiated cells could also elevate the total triglyceride content we established whether ADMA alters 3T3-L1 differentiation. Over the 72 hours of treatment from post-induction day 10 to 13 ADMA had no effect on the percentage of lipid laden cells (Supplementary Fig. 1a). Furthermore, ADMA treatment of 3T3-L1 cells throughout the entire differentiation period had no effect on the differentiation marker perilipin (Supplementary Fig. 1b). Sensitivity of fully differentiated 3T3-L1 cells to insulin was unaltered by ADMA treatment (Supplementary Fig. 1c). These observations are consistent with ADMA causing hypertrophy in 3T3-L1 cells predominantly via de-novo lipid synthesis.

### ADMA induces lipogenesis in HepG2 cells

As the liver is a significant site for ectopic fat deposition, we assessed whether ADMA treatment alters lipid accumulation in the human lipogenic hepatocyte-derived HepG2 cell line. ADMA increased the lipid content of HepG2 cells, as evidenced by increased BODIPY staining (Fig. 1e and f). As with 3T3-L1 cells SDMA and NOS inhibitors had no effect on lipid accumulation. HepG2 cell triglyceride content was increased by ADMA treatment (Fig. 1g). These data indicate that the effects of ADMA are seen in two of the most highly lipogenic cell types and in both mouse and human cells.

### ADMA stimulates mTOR driven lipogenesis

To investigate the effect of ADMA on de-novo lipid synthesis we assessed the involvement of sterol regulatory element-binding protein-1 (SREBP-1) signalling. ADMA (10 μM) treatment of 3T3-L1 cells for 6h increased SREBP-1 nuclear accumulation (Fig. 2a). Downstream of SREBP-1 ADMA increased the expression of its target genes acetyl-CoA carboxylase (*Acaca*), fatty acid synthase (*Fasn*) and HMG-CoA reductase (*Hmgcr*) (Fig. 2b). As mTOR (mammalian target of rapamycin) is important for SREBP-1 cleavage (Porstmann *et al*., 2009) we measured the expression and activity of this enzyme. ADMA induced the expression of mTOR at both the mRNA (Fig. 2c) and protein levels (Fig. 2d and e); an effect that was not reproduced by SDMA or NOS inhibition. The induction of mTOR was inhibited by actinomycin D (5ug/ml) treatment over 30 hours (Fig. 2f) confirming that ADMA induces transcription rather than altered mRNA stability. To confirm that upregulation of mTOR expression resulted in increased activity we performed western blotting for the mTOR target AKT Ser^473^ (Fig. 2g and h) which was upregulated by 1 and 3 μM ADMA. SREBP-1 cleavage and lipogenesis rely on mTORC1 signalling while AKT phosphorylation is mTORC2 dependent (Kim and Guan, 2019); therefore, long-term rapamycin treatment was utilised to effectively block both pathways and determine the effect this had on ADMA driven lipogenesis (Sarbassov et al. 2006). Treatment of 3T3-L1 cells for 48 hours with rapamycin (10 nM) blocked the ADMA induced cellular hypertrophy and the induction of *Acaca* expression (Fig. 2i and j). The NO donors S-nitroso-N-Acetyl-D, L-penicillamine (SNAP) as well as the stable cGMP analogue 8-bromo-cGMP had no impact on mTOR expression (Fig. 2k).

**Fig. 2:**
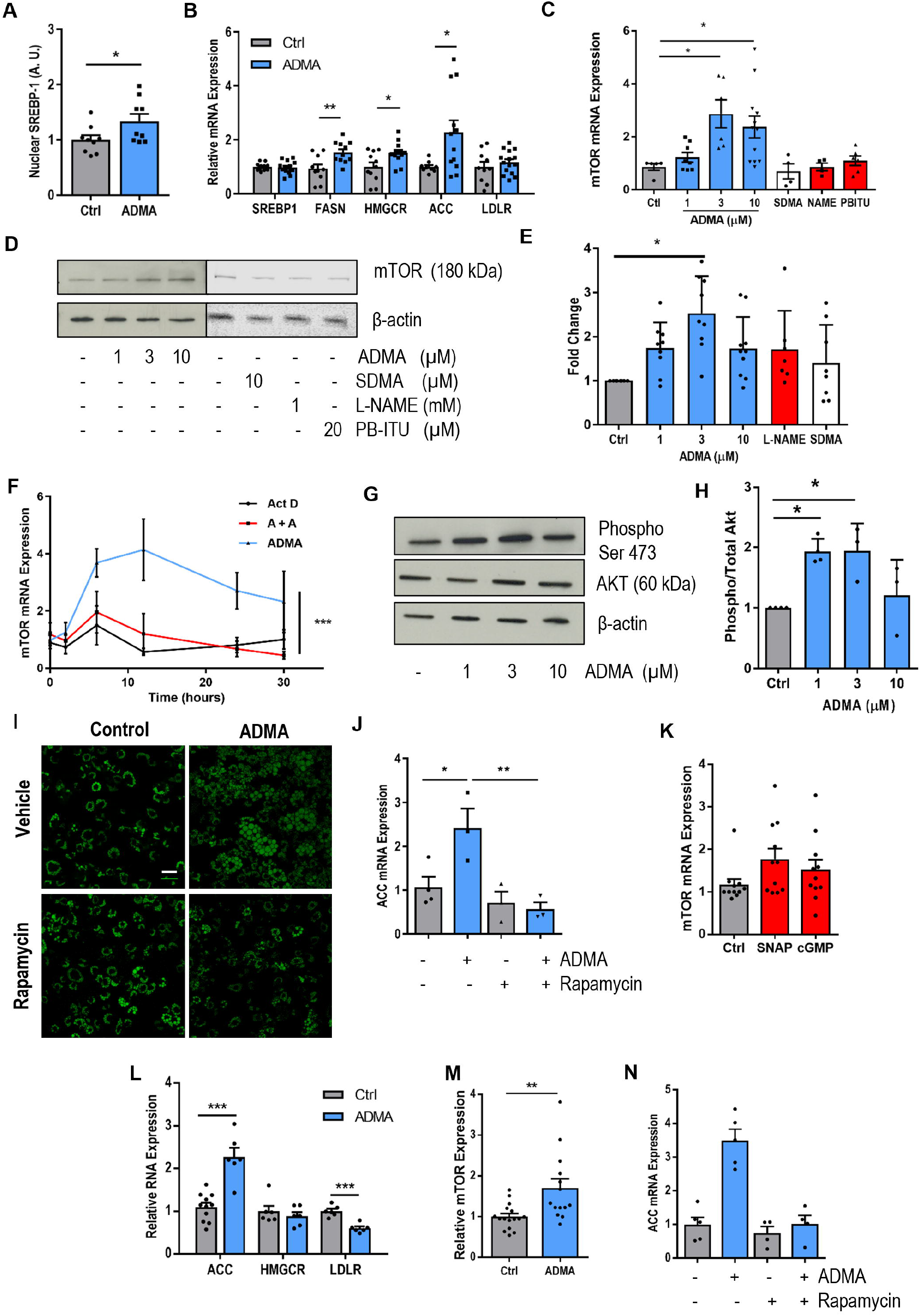
ADMA upregulates the mTOR-SREBP1 signalling pathway in 3T3-L1 and HepG2 cells cultured in DMEM. **(a)** Nuclear SREBP-1 levels in 3T3-L1 adipocytes following 6 hours stimulation with 10 µM ADMA (N=9). **(b)** qPCR analysis of SREBP target genes after stimulation of 3T3-L1 cells with 3 µM ADMA for 12 hours, values are fold change normalised against 18S housekeeper gene (N=12). **(c)** qPCR analysis of mTOR mRNA expression (N=5) and **(d and e)** western blot analysis of mTOR levels following 48 hour incubation of 3T3-L1 cells with ADMA, 10 µM SDMA, 1 mM L-NAME and 20 µM PBI-TU. Actin was used as a loading control, representative immunoblots are shown, densitometry analysis of 7 independent experiments. **(f)** qPCR analysis of a 30 hour timecourse with 10 µM ADMA in 3T3-L1 cells in the presence or absence of 5 µg/ml actinomycin D (Act D) (N=5, A + A = ADMA + Act D). **(g and h)** Western blot analysis of phosphorylation of AKT at Ser^473^ following 48 hours treatment, actin was used as a loading control, densitometry analysis of 4 independent experiments. **(i)** BODIPY staining of 3T3-L1 cells incubated with 10 µM ADMA with or without rapamycin (10 nM) for 72 hours, representative images from 3 independent experiments, scale bar 50 µm. **(j)** qPCR analysis of ACC expression after incubation of 3T3-L1 cells with ADMA (48 hours, 10 µM) in the presence or absence of 10 nM rapamycin (N=3). **(k)** qPCR of mTOR expression in 3T3-L1 adipocytes after 48 hours in the presence of 100 µM SNAP or 100 µM 8-bromo-cGMP (N=10). **(l)** qPCR analysis of lipogenesis gene mRNA expression in HepG2 cells incubated for 18 hours with 3 µM ADMA, normalised against 18S expression. (N=6) **(m)** qPCR analysis of mTOR expression in HepG2 cells after incubated for 18 hours in 3 µM ADMA (N=12). **(n)** qPCR analysis of ACC expression in HepG2 cells incubated for 18 hours in 3 µM ADMA in the presence or absence of rapamycin (10 nM). All qPCR are expressed as fold change normalised against 18S expression. Analysis was by one-way ANOVA followed by multiple comparisons test (Bonferroni) in **c, e, h, j, k** and **n**; a two-tailed students t-test in **a, b, l** and **m**, and a two-way ANOVA in **f**. * P<0.05, **P<0.01, ***P<0.001. Data presented are mean ± S.E.M.

We again utilised HepG2 cells to assess whether ADMA-mTOR signalling occurs in multiple cells types. ADMA again increased the transcription of SREBP-1 target genes *Acaca* and Ldlr (Fig. 2l) as well as mTOR (Fig. 2m). In HepG2 cells rapamycin also blocked the ADMA driven upregulation of *Acaca* (Fig. 2n). These data indicate that ADMA drives neo-lipogenesis through the upregulation of mTOR signalling and SREBP-1 activation in both adipocytes and hepatocytes.

### Adipocyte-specific DDAH1 deletion increases body weight and fat mass in mice

*In vivo*, dimethylarginine dimethylaminohydrolase (DDAH) enzymes metabolise 80% of asymmetric methylarginines with the remaining 20% excreted by the kidneys (Leiper *et al*., 2007). In 3T3-L1 cells we established DDAH1 was strongly upregulated through differentiation to mature adipocytes while DDAH2 was significantly down-regulated before returning to basal levels (Supplementary Figure 2a and b). Therefore, to elevate ADMA concentrations in mature adipocytes, we choose to develop a mouse in which DDAH1 was specifically deleted in mature adipocytes (termed DDAH1^Ad-/-^) using the adiponectin-BAC cre driver (Eguchi *et al*., 2011). DDAH1 floxed mice (DDAH1^fl/fl^) were used as controls (Dowsett *et al*. 2015). DDAH1 protein expression was reduced by approximately 75% in primary adipocytes (Supplementary Fig. 3a and b). DDAH2 expression was unaffected by DDAH1 deletion (Supplementary Fig. 3a and c). The ADMA content in primary epididymal adipocyte lysates was towards the limit of detection by mass spectrometry. ADMA in DDAH1^Ad-/-^ tended to be higher, but this did not reach statistical significance (Fig. 3a). However, adipocyte NOx content were significantly lower in DDAH1^Ad-/-^ mice (Fig. 3b) suggesting that there is a physiologically significant increase in ADMA. On the other hand, plasma concentrations of both ADMA and NOx were unaffected by DDAH1 adipocyte deletion (Fig. 3c and d). This is in keeping with other tissue specific DDAH1 knockout mice where systemic ADMA remains unchanged (Dowsett *et al*., 2015; Tomlinson *et al*., 2015). Male DDAH1^Ad-/-^ mice maintained on a normal chow diet weighed significantly more than littermate controls over a period of 6 to 22 weeks of age (Fig. 3e). This was due to increased fat mass (Fig. 3f) with no difference in lean mass (Fig. 3g). At 22 weeks of age DDAH1^Ad-/-^ mice had larger epididymal fat depots (Fig. 3h), and elevated plasma free fatty acids (Fig. 3i). Staining of epididymal fat pads demonstrated adipocyte hypertrophy in DDAH1^Ad-/-^ (Fig. 3j and k) which was accompanied by an increase in macrophage infiltration as detected by galectin-3 staining (Fig. 3j and l). Analysis of primary adipocytes isolated from DDAH1^Ad-/-^ mice indicated that both mTOR and ACC mRNA expression was significantly elevated compared to littermate controls (Fig. 3m) suggesting ADMA-induced mTOR signalling and adipocyte lipogenesis also occurs *in vivo*.

**Fig. 3:**
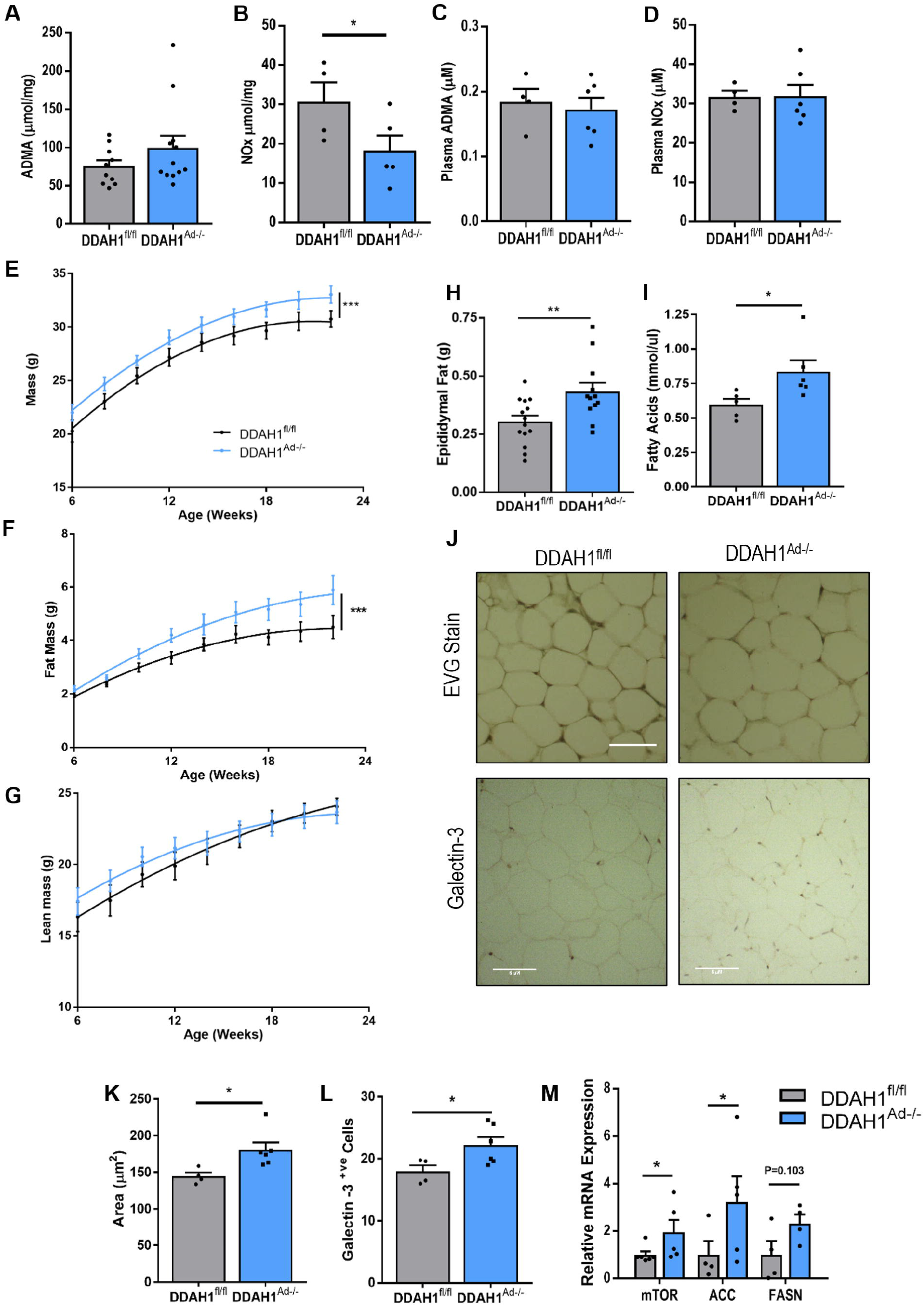
Deletion of ADMA metabolising enzyme DDAH1 in adipocytes increases body fat and adipocyte hypertrophy. DDAH1^Ad-/-^ and DDAH1^fl/fl^ littermate controls were assessed fortnightly from 6-22 weeks of age. At 22 weeks 1° adipocytes were isolated from epididymal fat pads to assess intracellular **(a)** ADMA (N=10) and **(b)** NOx (N=5). Plasma was assessed for circulating **(c)** ADMA and **(d)** NOx (N=4-5). DDAH1^Ad-/-^ and DDAH1^fl/fl^ **(e)** weight, **(f)** fat mass and **(g)** lean mass from 6 to 22 weeks of age. N=11-12 individual mice per group. **(h)** Epididymal fat mass, N=12-14; and **(i)** Plasma free fatty acids, N=6 at 22 weeks. **(j)** Epididymal fat was stained with EVG dye and Galectin-3 antibody to assess **(k)** adipocyte size and **(l)** macrophage infiltration (scale bar 50 µm, representative images with staining performed on 5 mice per group). **(m)** qPCR analysis of mTOR and ACC mRNA expression in primary adipocytes isolated from epididymal fat pads taken at 22 weeks of age, N=5 for each group. Analysis was by non-linear regression in **e-g;** a two-tailed students t-test in **a-d** and **h-m**. * P<0.05, **P<0.01, ***P<0.001. Data presented ± S.E.M.

To confirm that changes in adipocyte and adipose size in DDAH1^Ad-/-^ mice were due to changes in adipocyte signalling rather than a shift in whole body metabolism we utilised the CLAMS (Comprehensive Lab Animal Monitoring) system. VCO_2_ and VO_2_ were unaltered in DDAH1^Ad-/-^ (Supplementary Fig. 4a and b) and over the time period studied both groups consumed the same amount of food (Supplementary Fig. 4c). Surprisingly, DDAH1^Ad-/-^ mice showed a small but statistically significant increase in physical activity (Supplementary Fig. 4d) with higher total ambulatory counts. The mechanism underlying this change in activity is unclear and will require further investigation. These data are consistent with ADMA driving adipocyte hypertrophy and adipose expansion through a cell autonomous manner rather than through altered metabolic rate.

### Adipose ADMA is significantly raised by high fat feeding

In human subjects it has previously been reported that plasma ADMA concentrations are elevated in obese individuals (Eid *et al*., 2004; Kocak *et al*., 2011). To discover whether adipocyte DDAH1 plays a role in this increase we placed male DDAH1^Ad-/-^ and control littermates on a high fat diet (HFD) consisting of 60% calories from fat from 6 weeks old for a period of 16 weeks. Following high fat feeding both DDAH1^Ad-/-^ and DDAH1^fl/fl^ mice gained weight, due to increased fat mass with no change in lean mass compared to control mice on a normal chow diet (Supplementary Figure 5a-c). Interestingly, there was no difference between genotypes. Both DDAH^Ad-/-^ and Wt mice had enlarged epididymal fat pads and displayed extensive adipocyte hypertrophy (Supplementary Figure 5d and e). Unlike human studies there was no change in either plasma ADMA or NOx concentration in either strain following high fat feeding (Supplementary figure 5f and g). However, ADMA concentrations were very significantly increased in adipocyte lysates from both HFD-fed control and DDAH1^Ad-/-^ mice compared to mice on a normal chow diet, (Supplementary Figure 5h and i) suggesting that ADMA formation is of greater physiological importance than ADMA metabolism in obesity.

### Adipocyte DDAH1 expression is reduced in eNOS^-/-^ mice

To determine whether the effects of DDAH1^Ad-/-^ deletion *in vivo* is NO independent we investigated eNOS knockout mice over the same time period. As has previously been described eNOS^-/-^ had increased fat and lean mass at 22 weeks of age (Supplementary Figure 6a-c) and had significantly larger epididymal fat pads (Supplementary Figure 6d) (Nakata *et al*. 2008). Interestingly, DDAH1 expression was significantly suppressed in isolated adipocytes corresponding to an increase in adipocyte mTOR expression, suggesting ADMA may upregulate this pathway in eNOS knockout mice (Supplementary Figure 6e-g).

### ADMA induced lipogenesis via CaSR activation in 3T3-L1 cells

In comparison to the concentrations of ADMA required for NOS inhibition in cell culture systems (100 μM) the concentration of ADMA capable of driving adipocyte hypertrophy is low (1 µM). At this concentration ADMA entry into cells via the cationic amino acid transporter is likely to be limited due to competition from the high concentration of cationic amino acids in the culture media (Strobel *et al*., 2012). Therefore, we hypothesised that ADMA may act *via* an extracellular receptor. The calcium-sensing receptor (CaSR) is a member of the class C family of amino acid sensitive GPCRs (Conigrave and Hampson, 2010; Chun, Zhang and Liu, 2012). CaSR is expressed in both adipocytes and hepatocytes (Cifuentes *et al*., 2010) and has been linked to adipose dysfunction and fat accumulation (Bravo-Sagua *et al*., 2016; Villarroel *et al*., 2016). We confirmed CaSR expression within our 3T3-L1 and HepG2 cultures (Supplementary Figure 7) and that CaSR expression was unaffected by 3T3-L1 differentiation (Fig. 4a). Activation of CaSR with the positive modulator cinacalcet achieved a similar level of adipocyte hypertrophy and lipid accumulation as 10 µM ADMA (Fig. 4b-d). We then investigated whether inhibition of CaSR blocked the lipogenic effects of ADMA on 3T3-L1 cells. The CaSR inhibitors Calhex-231 (10µM) and NPS-2143 (10µM) inhibited the lipogenic effect of ADMA on both adipocyte cell size and lipid area. Interestingly, NPS-2143 alone but not Calhex-231 caused an increase in 3T3-L1 cell size (Fig. 4e-g); but both compounds antagonised the effect of ADMA. NPS-2143 inhibited ADMA induced lipid accumulation in HepG2 cells confirming CaSR signalling was not restricted to the adipocyte model (Fig.4h and i).

**Fig. 4:**
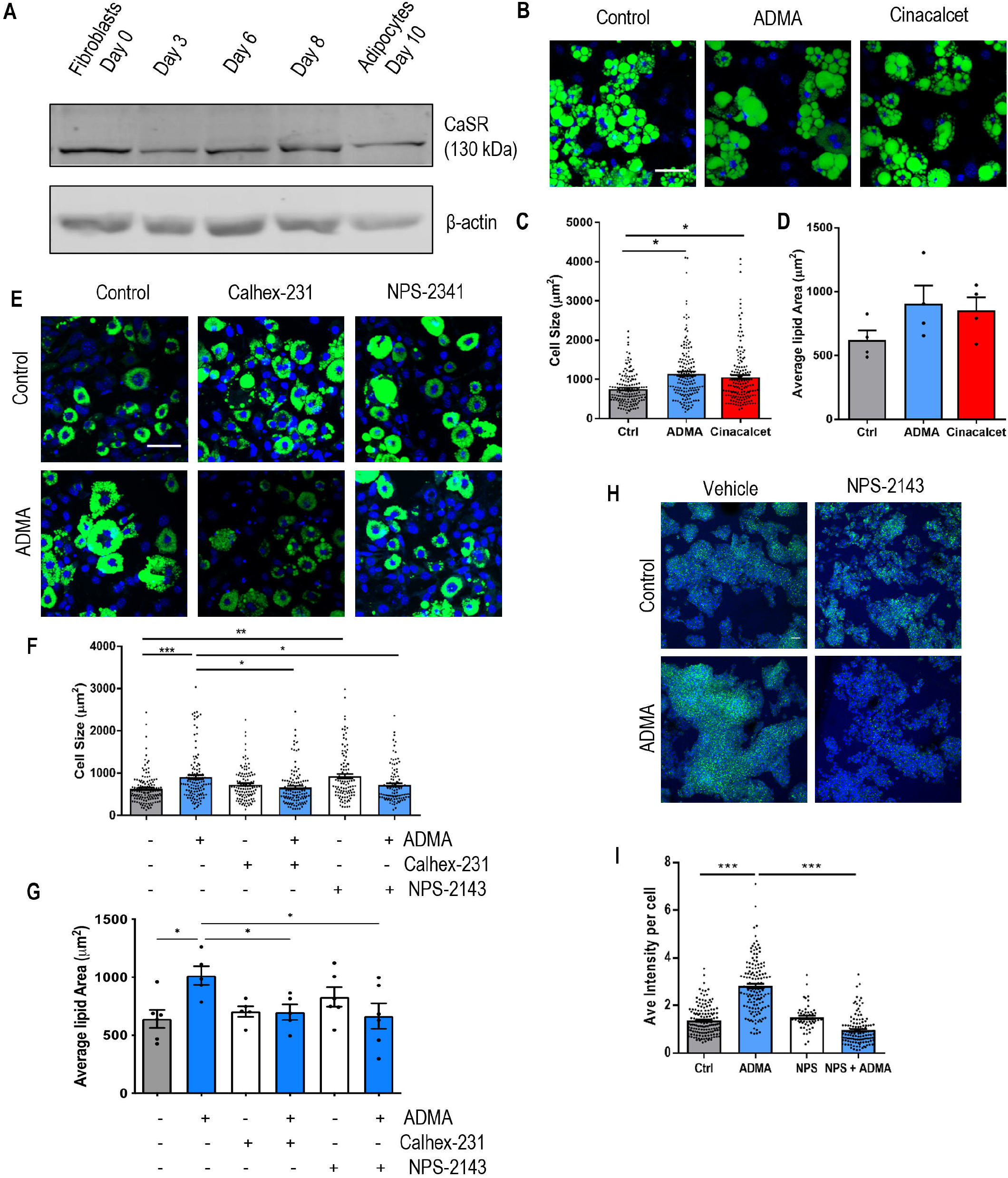
ADMA driven lipid accumulation is mimicked by CaSR agonists and blocked by CaSR antagonists when cultured in DMEM. **(a**) Western blot analysis confirms CaSR expression is unaffected by 3T3-L1 differentiation. **(b-d)** 3T3-L1 cells incubated for 72 hours with 10 µM ADMA or 50 nM Cinacalcet were **(b)** stained with BODIPY, representative images from 4 independent experiments, scale bar 50 µm and measured for **(c)** cell size and **(d)** average lipid area (N=160 cells from 4 independent experiments).**(e-g)** 3T3-L1 incubated for 72 hours with 10 µM ADMA in the presence of 10 µM Calhex-231 or NPS-2413 were **(e)** stained with BODIPY, representative images from 3 independent experiments, scale bar 50 µm and measured for **(f)** cell size (N=130 cells from 3 independent experiments) and **(g)** average lipid area (N=5). **(h)** Representative images from HepG2 cells treated with 10 µM ADMA in the presence of 10 µM NPS-2431 for 18 hours. **(i)** Lipid accumulation in HepG2 cells was established through average fluorescent intensity of BODIPY staining per cell. Analysis was by on-way ANOVA followed by Bonferroni multiple comparisons test in **c-d** and **f-g** and **i**. *P<0.05, **P<0.01, ***P<0.001. Data presented ±S.E.M.

### ADMA increases CaSR sensitivity

CaSR is a promiscuous GPCR signalling through G_q –_ resulting in increased cytosolic Ca^2+^, G_i/o_ inhibiting cAMP signalling and finally G_12/13_ Rho signalling (Conigrave and Ward (2013)). To determine if ADMA directly modifies CaSR activity we developed a HEK293-CaSR cell line using the Flp-In T-REX transfection system (Ward *et al*., 2011) which overexpresses CaSR following doxycycline treatment (Fig. 5a and b). To assess whether ADMA affects intracellular Ca^2+^ mobilization we used the potent CaSR agonist Gd^3+^ to increase [Ca^2+^]_i_ in a dose-dependent manner. Gd^3+^ and Cinacalcet did not stimulate intracellular Ca^2+^ mobilisation in vector only cells or HEK-CaSR cells untreated with doxycycline (data not shown). ADMA increased CaSR sensitivity to Gd^3+^ (Fig. 5c and d) shifting the dose-response curve to the left (Control EC_50_ - 0.2 mM, ADMA EC_50_ - 0.06 mM) and to a similar degree as previously reported for other amino acids (Geng *et al*., 2016; Zhang *et al*., 2016). Pre-treatment with NPS-2143, a CaSR inhibitor, completely inhibited the effect of ADMA on intracellular Ca^2+^ (Fig. 5e). As L-arginine competes with ADMA for NOS binding we investigated whether the same is true for CaSR. Increasing L-arginine to physiological (80 µM) or supra-physiological (1 mM) concentrations had no effect on CaSR signalling either basally or in the presence of ADMA (Fig. 5f). CaSR is also known to regulate cAMP concentrations via G_i/o_ signalling. Basally neither Gd^3+^ nor ADMA altered cAMP concentrations; however, following the addition of forskolin to induce adenylyl cyclase activity Gd^3+^ inhibited cAMP production, an effect that was enhanced in the presence of ADMA (Fig. 5g). As Gd^3+^ is not an endogenous agonist for CaSR we confirmed that ADMA also had an effect on extracellular Ca^2+^ stimulated CaSR activation. Using intracellular Ca^2+^ mobilisation to visualise CaSR activation ADMA (10 μM) significantly left-shifted the dose-response curve to a similar degree as the known CaSR modulator phenylalanine (100 μM) compared to the PBS control (Figure 5h; EC_50_ - Control 1.7 mM Ca^2+^ ± 0.19, EC_50_ - Phenylalanine 0.76 mM Ca^2+^ ± 0.11, EC_50_ - ADMA 1.1 mM Ca^2+^ ± 0.10, P<0.05).

**Fig. 5:**
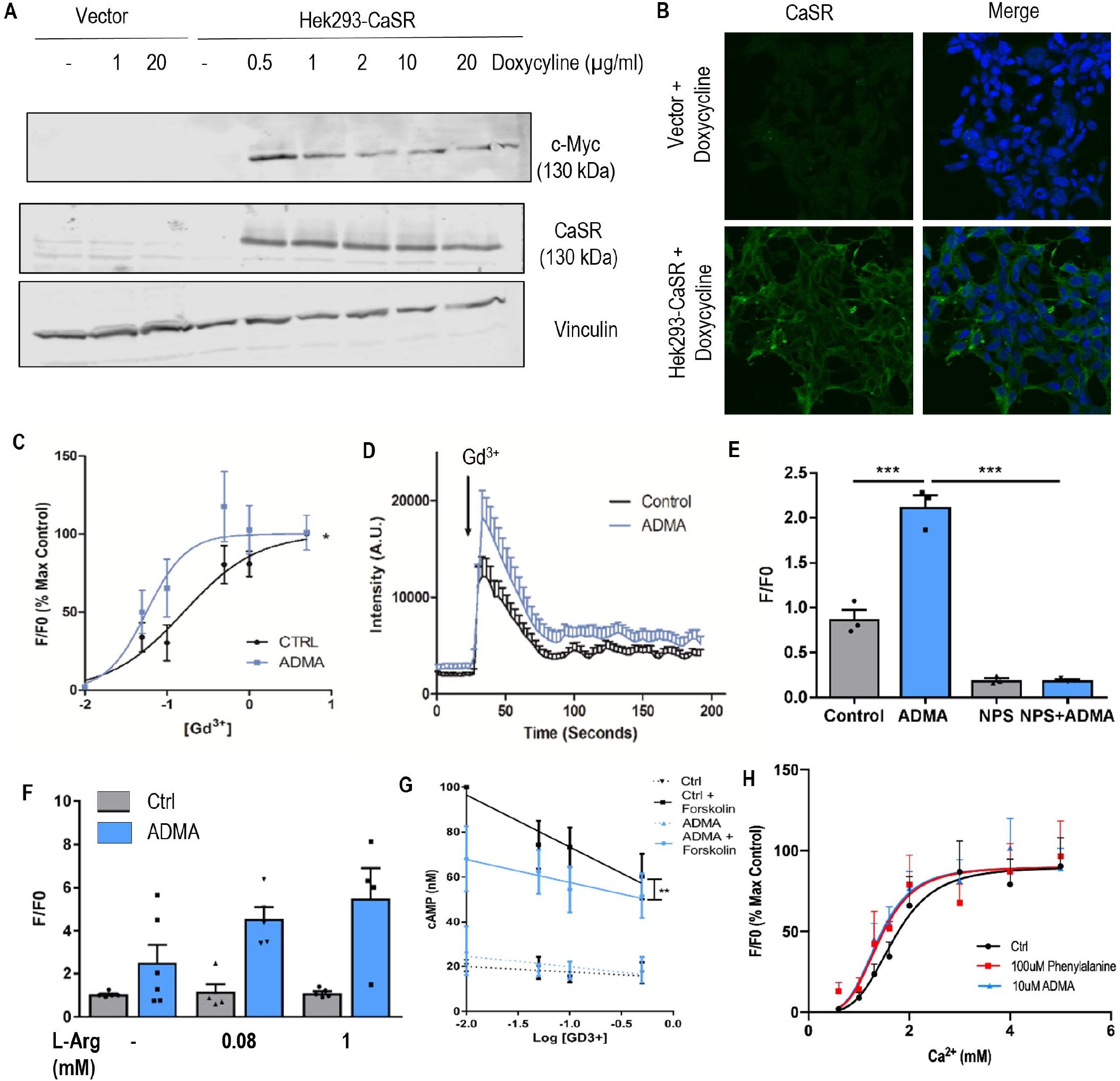
ADMA is a positive allosteric modulator for CaSR. HEK293-CaSR cells were cultured with doxycycline for 24 hours to stimulate CaSR expression as shown by **(a)** western blot and **(b)** immunostaining, representative images of 3 independent experiments. **(c-f)** HEK293-CaSR cells in a Ca^2+^-free salt solution were incubated with Cal520 Ca^2+^ sensitive dye and stimulated with increasing concentrations of Gd^3+^ alone or in the presence of 10 µM ADMA. **(c)** Dose-response curve, change in intensity from baseline normalised to the 100% response in control cells (N=12). **(d)** A representative trace of one experiment made from an average of 10 single cells analysed; 0.05 mM Gd^3+^ added at the arrow. **(e)** Cells were stimulated with 0.05 mM Gd^3+^ in the presence of 10 µM ADMA ± 10 µM NPS-2143 (N=3). **(f)** 0.05 mM Gd stimulation in the presence of varying arginine concentrations (N=5). **(g)** Intracellular cAMP concentration of HEK293-CaSR stimulated with increasing Gd^3+^ concentration ± 10 µM ADMA. Forskolin was used to stimulate cAMP production by adenylyl cyclase (Basal levels N=5, Forskolin stimulated N=8). **(h)** HEK293-CaSR incubated with Cal520 dye were cultured in 0.3 mM Ca^2**+**^ before being stimulated with increasing doses of Ca^2+^ in the presence of 100 µM Phenylalanine or 10 µM ADMA (N=6). Analysis was by a one-way ANOVA followed by multiple comparisons test (Bonferroni) in **e** and **f**; non-linear regression in **c, g and h**. * P<0.05, **P<0.01, ***P<0.001. Data presented mean ± S.E.M.

### SDMA may act as a competitive antagonist at the amino acid site on CaSR

As a range of amino acid species are known to bind CaSR and modulate its signalling, it is interesting that ADMA causes adipocyte hypertrophy whereas the methylarginine analogue SDMA does not (Figure 1a and b). Therefore, we modelled both ADMA and SDMA within the CaSR amino acid binging site. L-Tryptophan has previously been shown to bind CaSR (Geng *et al*., 2016; Zhang *et al*., 2016), the superimposition of ADMA within this site (Fig. 6a) shows both the amino acid moiety and guanidine are well accommodated. The alkyl sidechain makes close interactions with hydrophobic residues on the opposite side of the “hinge” region, particularly ILE416. The overlay of SDMA with ADMA (Fig. 6b) show that again the amino acid and guanidine are accommodated, however, the differing methylation state of SDMA impacts how the alkyl portion of the side chain can be accommodated. The close interactions between ADMA and ILE416 are not observed in docked SDMA due to the alternative positioning required to accommodate the symmetrical *N*^ω^-methylation. These results provide supporting evidence for L-ADMA binding at the L-Trp site and are consistent with our finding that ADMA can serve as a CaSR activator by making interactions that reinforce the closed conformation of CaSR, and that SDMA does not. Given that SDMA seems to be able to bind to CaSR but perhaps not actively modulate it we hypothesised that SDMA may act to antagonise positive allosteric binding. Incubation of HEK-CaSR cells with 100 μM phenylalanine increased intracellular Ca^2+^ mobilisation in response to stimulation with 1.6 mM Ca^2+^ (Figure 6c) as previously demonstrated in Figure 5h. Co-incubation with increasing SDMA concentrations (30-1000 μM) showed a dose-dependent trend to suppress the effect of phenylalanine. Finally, to determine whether SDMA has a physiological effect on CaSR activity we incubated differentiated 3T3-L1 cells with 10 µM ADMA in the absence or presence of excess (100 µM) SDMA (Fig. 6d and e). The presence of SDMA blocked ADMA driven adipocyte hypertrophy suggesting that the ADMA/SDMA ratio may be an important modulator of ADMA-CaSR signalling.

**Fig. 6:**
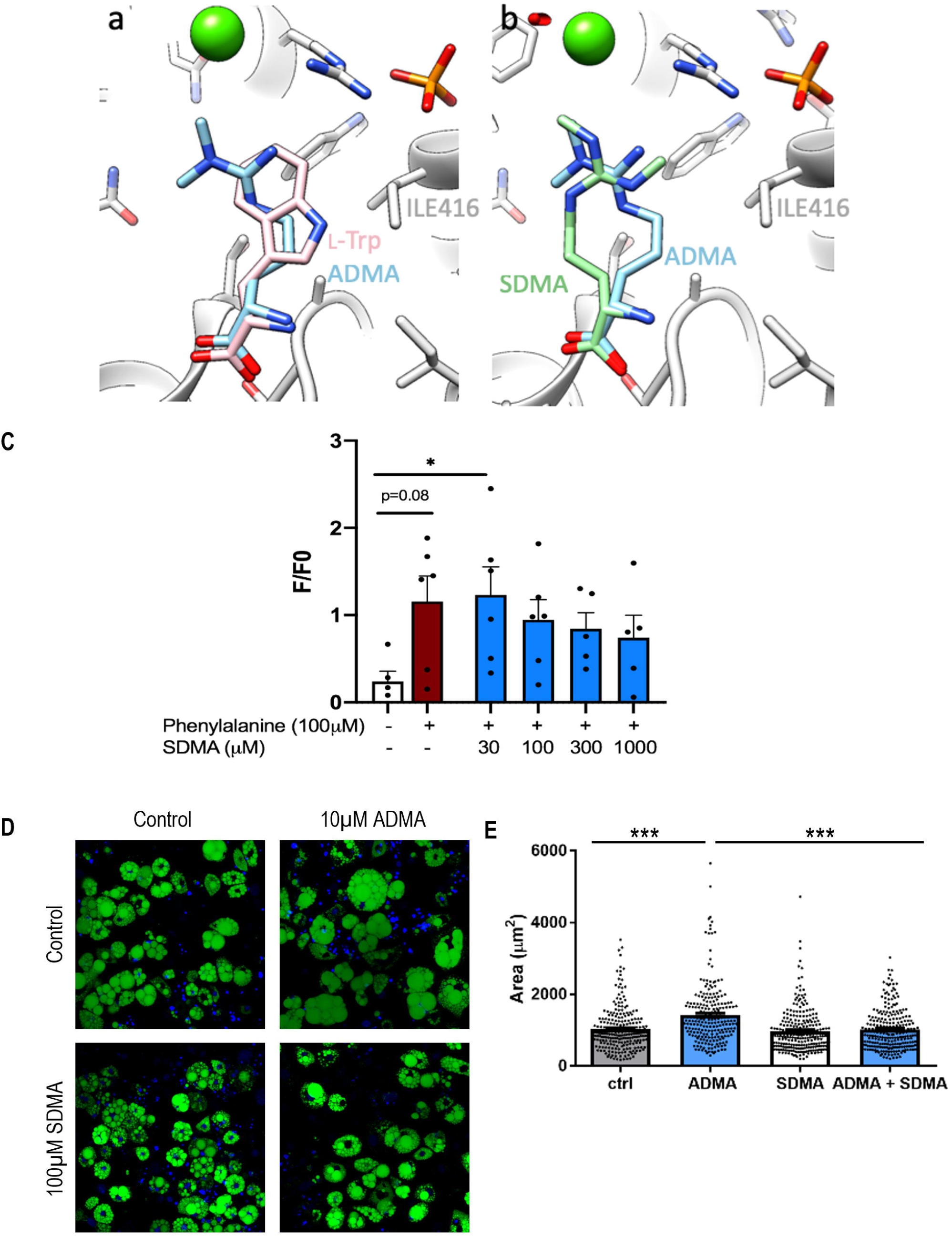
SDMA can act as a competitive inhibitor of ADMA. The active form of CaSR was modelled in Autodock Vina to predict the binding of ADMA in the L-Trp binding pocket **(a)** and then to compare the binding of SDMA with ADMA **(b). (c)** HEK293-CaSR cells were incubated with Cal520 Ca^2+^ sensitive dye and stimulated with 1.6 mM Ca^2+^ in the presence of 100 µM phenylalanine with or without increasing concentrations of SDMA (3-1000 µM) (N=6). 3T3-L1 cells were incubated for 72 hours with 10 µM ADMA in the presence of 100 µM SDMA **(d)** stained with BODIPY, (representative images from 4 independent experiments, and **(e)** measured for cell size. (From 4 independent experiments). Analysis by one-way ANOVA followed by Bonferroni multiple comparisons test in **c** and **e**. *P<0.05.

## Discussion

ADMA is an independent risk factor for cardiovascular disease with elevated plasma concentrations being associated with obesity in clinical studies (Eid *et al*., 2004; Kocak *et al*., 2011). However, an increase in the plasma concentration of many amino acids has been observed in obesity (Katsanos and Mandarino, 2011); therefore, we set out to establish whether ADMA is simply a marker of metabolic disease or a mediator of disease pathology and progression. Here we show for the first time that ADMA has a direct effect on adipocyte physiology and propose a new mechanism through which excess ADMA, or DDAH dysregulation, leads to adipocyte hypertrophy via stimulation of the calcium-sensing receptor (Fig. 7).

**Figure 7:**
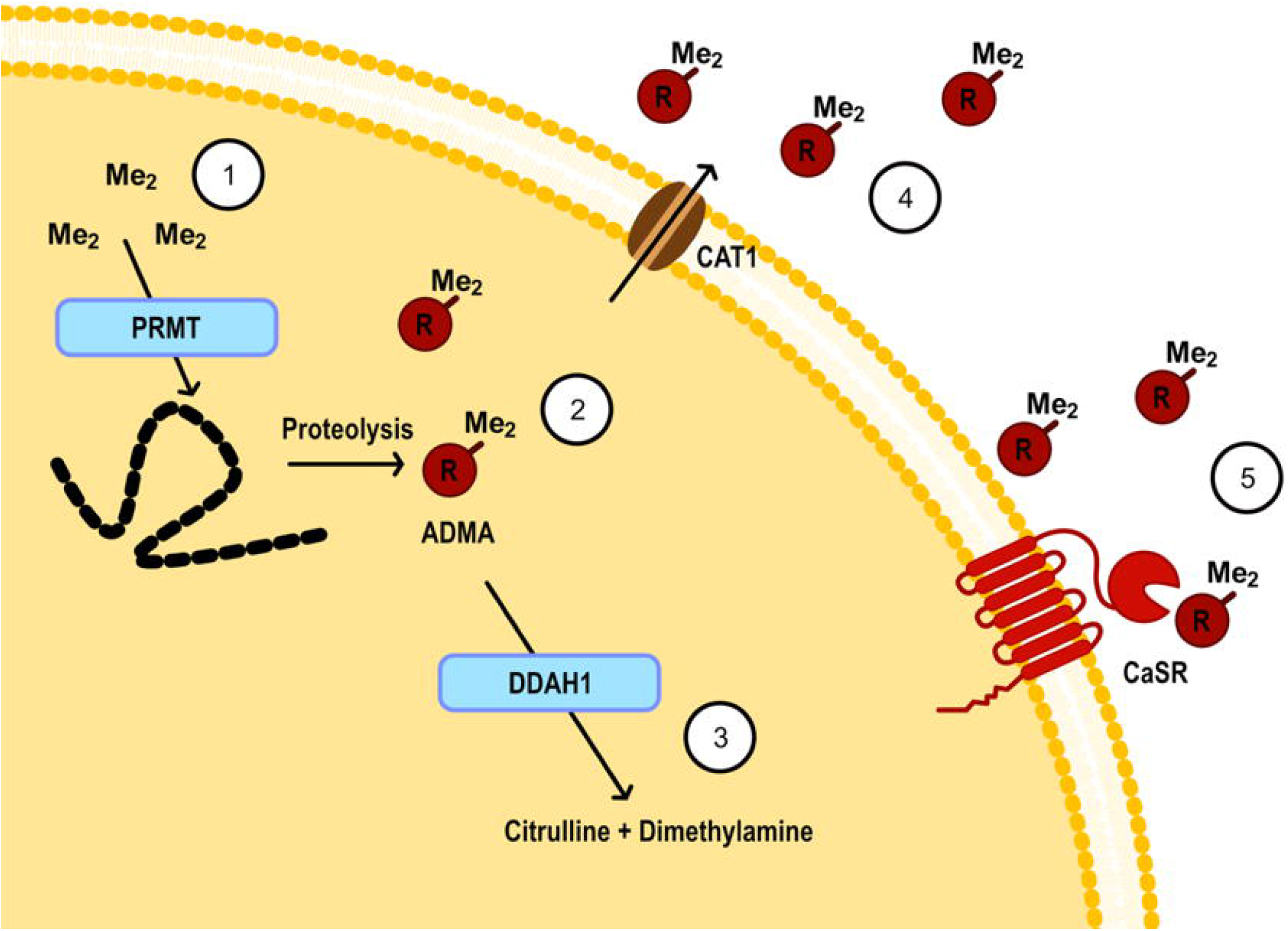
The ADMA-CaSR pathway. **(1)** Arginine residues within proteins can be methylated by the protein methyltransferase (PRMT) enzymes as one of many post-translational modifications. **(2)** There is no know demethylation step for functioning proteins therefore, following proteolysis, methylarginines including asymmetric dimethylarginine (ADMA) are released into the cytosol. **(3)** The major pathway for ADMA removal is via the dimethylarginine dimethylaminohydrolase (DDAH) enzymes which break down ADMA to citrulline and dimethylamine. **(4)** However, if DDAH is dysfunctional or there are high levels of ADMA production then ADMA can exit the cell through amino acid transporters, for example the cationic amino acid transporter (CAT1). **(5)** A build-up of ADMA in the extracellular space allows ADMA to bind to the calcium-sensing receptor (CaSR) as a positive allosteric modulator sensitising CaSR to its primary agonist Ca^2+^.

To examine what role ADMA plays in adipose physiology we employed the well characterised 3T3-L1 model, in which we established that prolonged (72 hour) exposure to ADMA increased cellular lipid content. In keeping with this, ADMA upregulated the mTOR-SREBP1 pathway leading to increased transcription of the lipogenesis genes *Fasn, Acaca* and *Hmgcr*. Similarly, HepG2 cells accumulated lipid when treated with ADMA suggesting this pathway is not exclusively restricted to adipocytes and may be a common feature among lipogenic cells.

To explore this pathway i*n vivo* we developed a novel mouse deficient in DDAH1 specifically in mature adipocytes. In these mice reduced adipocyte methylarginine metabolism was sufficient to increase visceral fat mass on a normal chow diet with a corresponding increase in mTOR and lipogenesis genes as seen in 3T3-L1 cells. The AdipoQ BAC cre utilised in this mouse model (Eguchi *et al*., 2011) removed the potential confounding effect of DDAH1 deletion in pre-adipocytes; although our experiments in 3T3-L1 cells would suggest that ADMA has no effect on adipocyte differentiation. In contrast when placed on a HFD differences between DDAH1^Ad-/-^ and DDAH1^fl/fl^ mice were lost (Supplementary Fig. 5); with the predominant effect a very significant increase in adipose tissue ADMA concentrations in both strains. This would suggest that elevated ADMA concentrations in obesity are a result of increased production rather than dysfunctional ADMA metabolism. Our data presented here contrasts with the observations of Li *et al*. (2017) who reported that mice globally deficient in DDAH1 gain more fat mass than wild type controls when both are fed a high fat diet. This may suggest that further increases in ADMA concentration including elevated plasma ADMA due to global loss of DDAH1 have a greater effect on adipocyte and hepatocyte function, not seen here due to the tissue specificity.

Although the DDAH1^Ad-/-^ mouse reflects the pathways identified in 3T3-L1 and HepG2 cells following ADMA treatment, DDAH1 deletion also resulted in a significant decrease in adipocyte NOx concentrations. Therefore, we cannot rule out that a reduction in NO production plays a role in fat accumulation in this mouse model. eNOS knockout mice have previously been shown to have increased visceral fat tissue and undergo adipocyte hypertrophy (Nakata *et al*. 2008) with decreased mitochondrial activity and impaired β-oxidation resulting in increased plasma free fatty acid levels (Gouill e*t al*., 2007). However, here we show that eNOS knockout mice (Supplementary Fig. 6) also have decreased adipocyte DDAH1 expression suggesting dysfunctional ADMA metabolism occurs in this mouse model as well as reduced NO production with both pathways possibly contributing to adipocyte hypertrophy and visceral obesity.

ADMA is a competitive inhibitor of all three NOS isoforms (Vallance *et al*., 1992a; Cardounel and Zweier, 2002). However, unexpectedly the synthetic NOS inhibitors L-NAME and PBI-TU did not replicate ADMA driven adipocyte hypertrophy or upregulation of mTOR expression in cell culture models. Our data indicates that ADMA has significant NO-independent biological effects at pathophysiological concentrations (1-3 µM). Cardounel *et al*. (2007) have calculated that these low concentrations would lead to at most a 10% inhibition of NOS. In contrast concentrations of ADMA that are capable of producing significant NOS blockade are only reached during the later stages of chronic kidney disease (Vallance *et al*., 1992a). Taken together these observations suggest a concentration dependent shift in the balance of ADMA signalling from NO-independent mechanisms towards direct NOS inhibition may play a significant role in the loss of cellular homeostasis that underlies human disease. Consistent with this hypothesis, elevation of plasma ADMA concentrations by on average 0.3 µM from ∼0.95µM to 1.3µM have been observed in metabolic disorders whereas increases of 1.0-4.0 µM are found in cardiovascular disease states associated with chronic renal failure.

Previous studies have suggested that ADMA may have actions in addition to NOS inhibition. In our laboratory we examined the effect of physiological and pathophysiological concentrations of ADMA on cultured endothelial cells. We identified ∼50 genes that were regulated in response to ADMA and demonstrated that for some of these genes the effect was not replicated by a synthetic NOS inhibitor (L-NIO) (Smith *et al*., 2005). The pathways regulated by ADMA included BMP and Osteocalcin signalling both of which have been shown to be regulated by CaSR. Our identification of a novel ADMA/CaSR signalling pathway now provides a plausible mechanism that might mediate these effects. Further studies will be required to elucidate the full range of ADMA/CaSR signalling in different cells and tissues. In addition to our own observations, Juretic and co-workers (Juretic *et al*., 1996) have reported that the induction of IL-2 in L-NMMA treated cultured PBMC’s is not replicated by treatment with synthetic inhibitors of NOS. Further work will be needed to establish whether L-NMMA can interact with and modulation CaSR activity.

Our data for the first time identifies a NO-independent receptor for ADMA, the amino acid sensitive GPCR CaSR. CaSR has been previously implicated in lipid homeostasis and its expression is upregulated in fully differentiated adipocytes (He *et al*., 2011). The CaSR agonist GdCl_3_ increases lipid accumulation and pro-adipogenic gene expression in the SW872 pre-adipocyte cell line (He *et al*., 2011); while cinacalcet increases HepG2 triglyceride content in culture (Villarroel *et al*., 2016). Furthermore, Rybchyn et al. (2019) have recently demonstrated CaSR-mediated activation of mTOR complex 2 signalling leading to increased phosphorylation of AKT. Interestingly, this study highlights the importance of the scaffolding protein homer-1 as a key player in linking CaSR to mTORC2 signalling. Currently, our hypothesis built on our modelling studies is that ADMA and SDMA bind directly to the amino acid site of CaSR and alter activity. However, further studies will be necessary to fully understand whether ADMA alters CaSR activity by altering CaSR protein complexes.

Whilst ADMA competes with L-arginine for binding to the active site of NOS our data presented here indicates that arginine is unable to compete with ADMA for CaSR. The literature relating to L-arginine binding to CaSR is somewhat contradictory with some studies reporting no effect of basic amino acids on the receptor (Conigrave and Ward, 2013) while others report a strong stimulation of CaSR by L-arginine in the gut (Mace, Schindler and Patel, 2012). In our hands, using stably transfected HEK293 cells, L-arginine was unable to directly activate CaSR or modulate the effect of ADMA on the receptor. However, these experiments were performed in a setting without calcium present in the buffer, and previous studies have shown that L-amino acids in general have a greater effect on CaSR activation at higher (2.5 mM) Ca^2+^ concentrations (Conigrave et al., 2000). L-arginine in particular, seems sensitive to the prevailing Ca^2+^ concentration only showing a low potency to activate CaSR at around 2mM Ca^2+^ and therefore, the absence of calcium here may mask the full effect of L-arginine on CaSR activity (Conigrave et al., 2004). A full series of experiments exploring L-arginine and ADMA interacting at the CaSR will need to be performed at a range of Ca^2+^, L-arginine and ADMA concentrations. ADMA increased CaSR sensitivity to Gd^3+^ and Ca^2+^ in its physiological range (low µM) suggesting greater affinity for CaSR than the known CaSR modulators L-phenylalanine and L-tryptophan both of which have been shown to act in the mM range (Zhang *et al*., 2016). In contrast to the lack of effect of L-arginine in our studies we have demonstrated that SDMA is able to block the effect of ADMA and phenylalanine on CaSR. These observations are of potential physiological importance as elevated SDMA concentrations are significantly associated with cardiovascular risk but, to date, no mechanism of action that can explain this association has been identified. Our *in silico* modelling studies suggest that both ADMA and SDMA can bind in the amino acid binding pocket of CaSR that crystallographic studies have demonstrated is occupied by tryptophan. Both methylated arginine molecules are predicted to bind with equal affinity however only ADMA is predicted to interact with Ile416 in the activation loop of CaSR. These observations provide a potential molecular explanation of the effects of ADMA/SDMA on CaSR activity. Structural studies will be required to fully understand the structure activity relationships of ADMA/SDMA:CaSR and the impact of genetic variants in CaSR on methylarginine signalling via this receptor.

CaSR is a member of the family C amino acid-sensing GPCRs which are conserved in many species. The most closely related family C member to CaSR is GPRC6a which is activated directly by amino acids (particularly L-ornithine and L-arginine) and is modulated by Ca^2+^. This receptor has also been suggested to interact with methylarginines in the absence of Ca^2+^ albeit at supra-physiological concentrations (Christiansen *et al*., 2006) perhaps suggesting that ADMA-receptor signalling may be more widespread than CaSR alone. This proposition is supported by studies that have demonstrated that asymmetrically mono-methylated arginine (L-NMMA) can inhibit canavanine sensing by the Drosophila DmXR, a family C receptor related to mGluR (Mitri *et al*., 2009), suggesting that methylated arginine signalling *via* GPCRs may be an ancient and conserved mechanism.

Our identification of an ADMA-CaSR signalling pathway that is sensitive to small changes in ADMA concentration in the micromolar range and is independent of prevailing arginine concentrations provides a potential mechanistic explanation of the association between ADMA and cardiovascular and metabolic risk, therefore offering novel therapeutic opportunities to mitigate the effects of elevated ADMA. Further studies will be required to fully understand the significance of ADMA signalling *via* GPCRs and the potential for this pathway to link catabolic and metabolic cellular processes *via* regulation of mTOR expression and activity.

## Acknowledgements

We would like to thank Prof. Graeme Milligan for the gift of the Flp-In T-REX cell line. This study was supported by MRC intramural funding to J.L and British Heart Foundation Centre of Research Excellence Award (RE/13/5/30177). E.H is a recipient of a British Heart Foundation 4 year studentship (FS/17/63/33485).

## Author Contributions

J.L conceived and managed the project. L.Dowsett, L.Duluc, E.H. F.A and I.S performed the *in vitro* experiments. L.Dowsett and O.B performed the animal experiments. L.Dowsett and L.Duluc performed the analysis. W.F. performed the protein modelling. L.Dowsett, L.Duluc and J.L wrote the manuscript with E.H and I.S providing critical appraisal of the manuscript. Correspondence and request for material should be addressed to J.L or to L.Dowsett

## Supplementary Materials

This report includes additional supplementary figures 1-7.

