## Supplementary Figure 1-7 for "Asymmetric dimethylarginine positively modulates Calcium-Sensing Receptor signalling to promote lipid accumulation and adiposity"

Supplementary Fig. 1

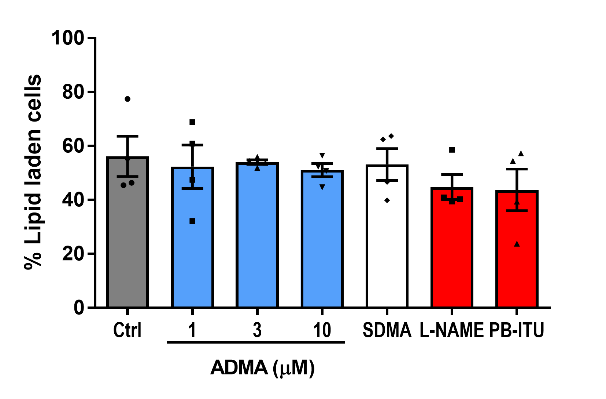

**a**

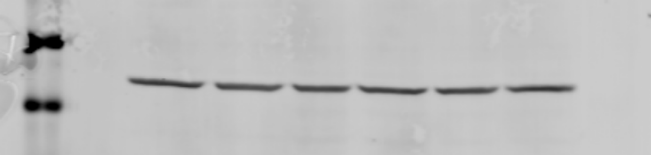

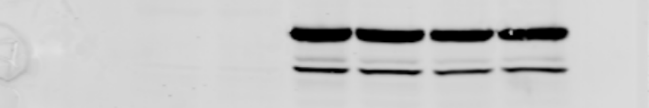

Perilipin-1

(65 kDa)

β-Actin

Fibroblasts

Adipocytes

Ctrl

ADMA

Ctrl

ADMA

SDMA

**b**

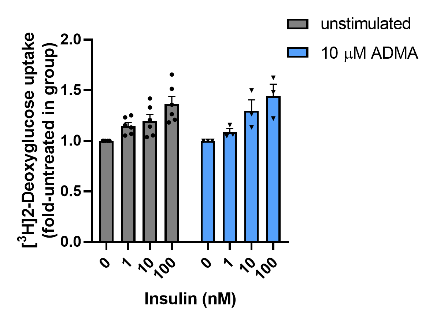

**c**

**Supplementary Fig. 1: ADMA has no effect on 3T3-L1 differentiation. (a)** The percentage of lipid laden cells determined by BODIPY and DAPI staining following 72 hour treatment by ADMA, 10 µM SDMA, 1 mM L-NAME and 20 µM PB-ITU. **(b)** Western blot of perilipin expression following 10 µM ADMA or SDMA treatment through the differentiation procedure. **(c)** Insulin sensitivity was assessed via insulin-stimulated uptake of [H^3^]-2-deoxyglucose after incubation in the presence or absence of 10 µM ADMA for 48 h.

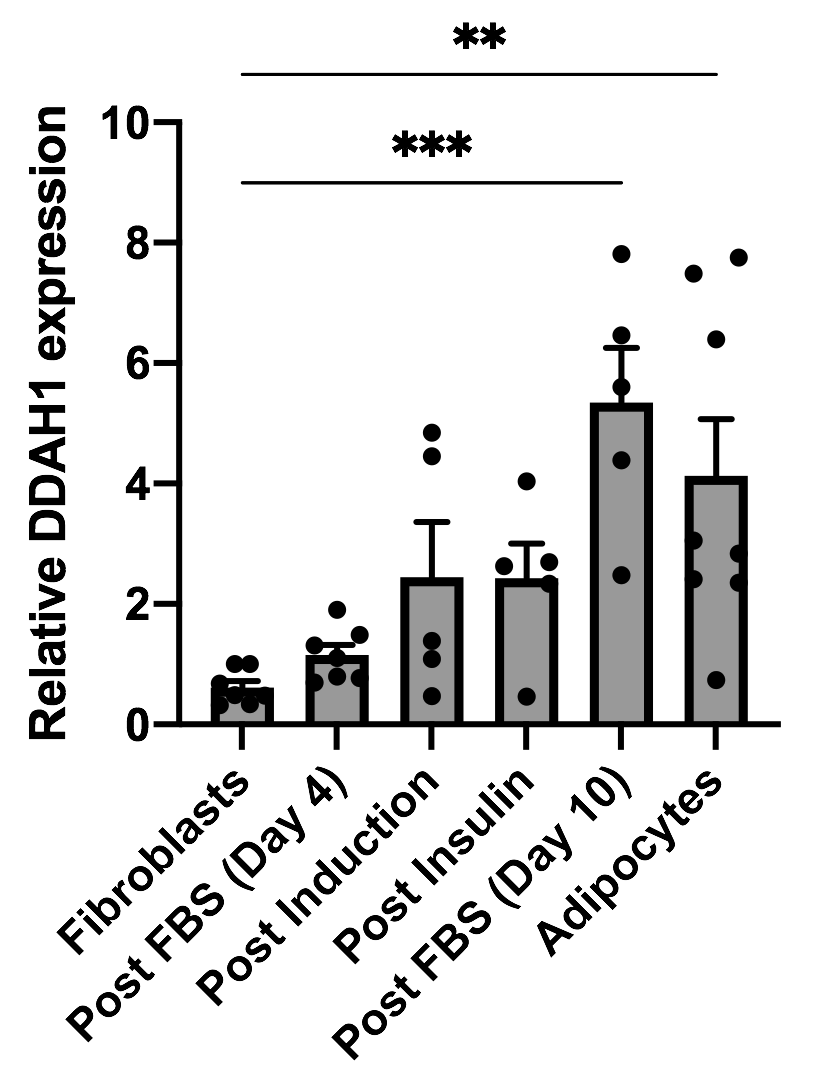

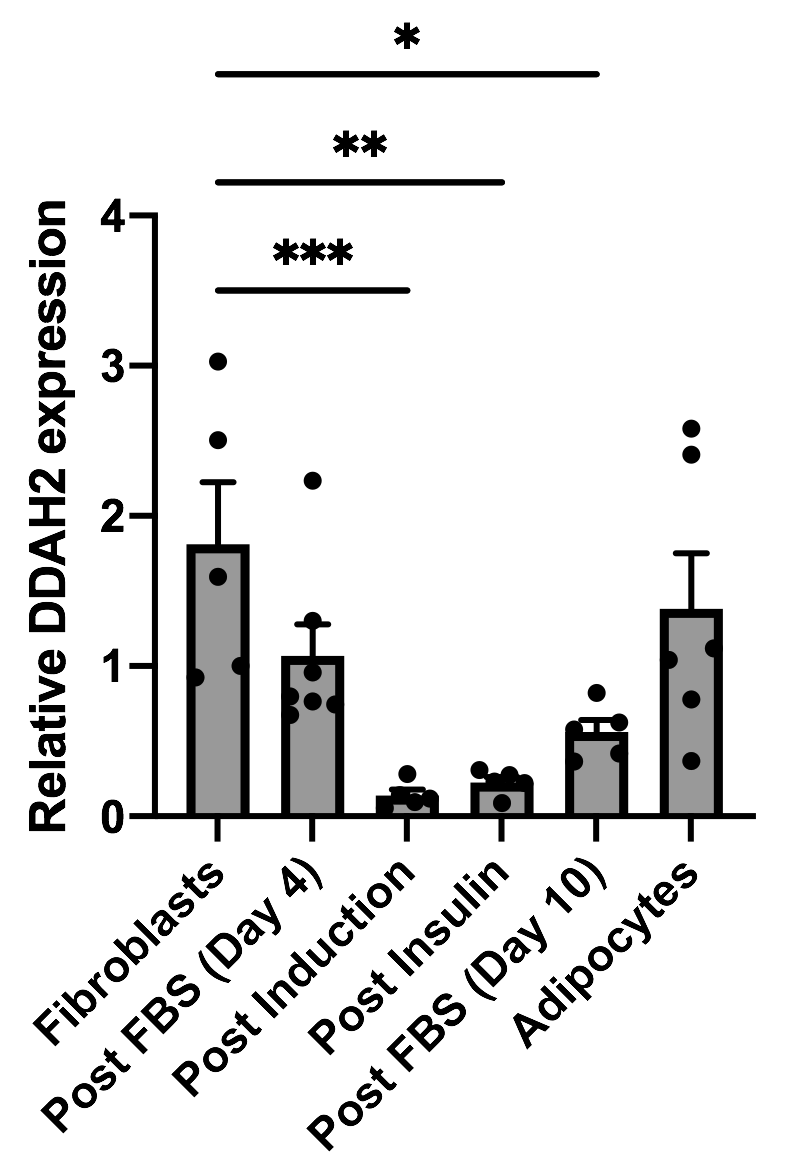

**Supplementary Fig 2: DDAH1 is strongly upregulated during 3T3-L1 differentiation.** Relative expression of **(a)** DDAH1 and **(b)** DDAH2 through 3T3-L1 differentiation determined by qPCR analysis. Analysis by one-way ANOVA followed by Bonferroni post hoc test. * p<0.05, **p<0.01, ***p<0.001

β-Actin

Supplementary Fig. 3

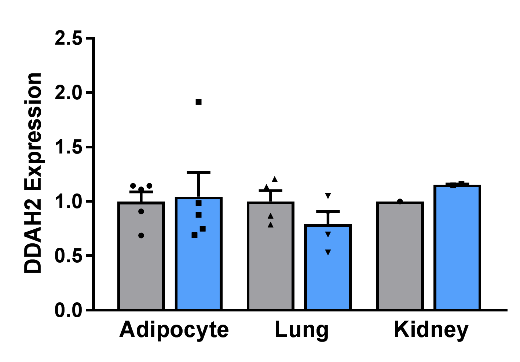

**a**

**b**

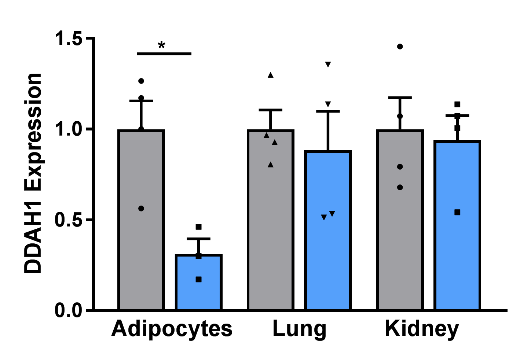

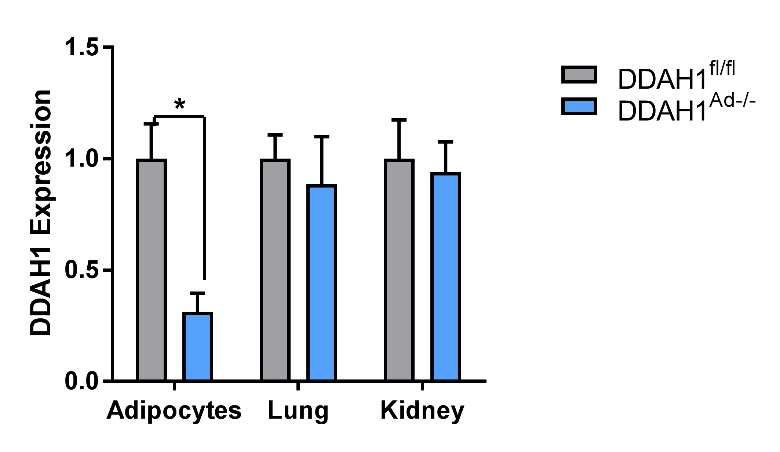

1° Adipocytes	Lung	Kidney

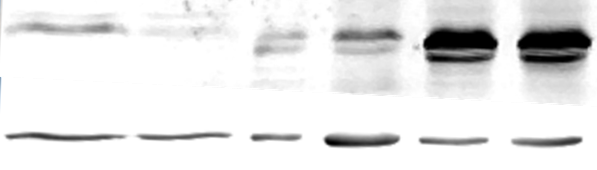

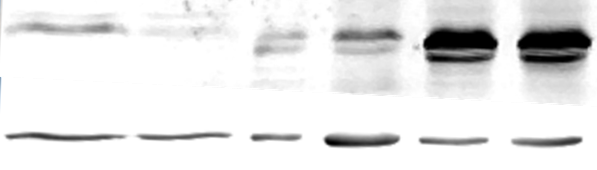

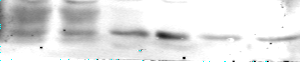

DDAH2

(32 kDa)

**+/+	-/-	+/+	-/-	+/+	-/-**

**c**

DDAH1

(38 kDA)

**Supplementary Fig. 3: DDAH1 is deleted in DDAH1^ad-/-^ adipocytes with no effect on DDAH2 expression. (a)** Western blot analysis of DDAH1 and DDAH2 expression in primary adipocytes, the lung and the kidney of DDAH1^Ad-/-^ and DDAH1^fl/fl^ mice. Representative image of 3 independent experiments. **(b and c)** Densitometry analysis of western blot data for DDAH1 and DDAH2. *P<0.05.

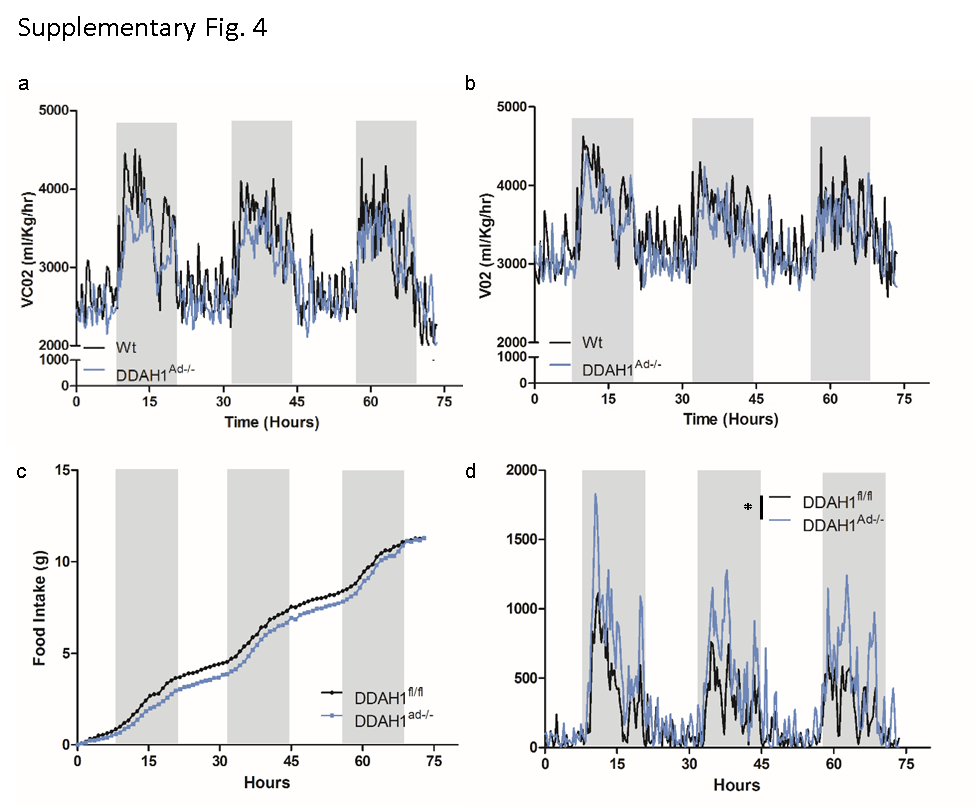

**Supplementary Fig.4: DDAH1^Ad-/-^ have unaltered metabolism and food intake but increased ambulatory activity.** DDAH1^fl/fl^ and DDAH1^Ad-/-^ were placed in a Comprehensive Lab Animal Monitoring Systems (CLAMS) to assess whole body calorimetry over a 72 hour period. **(a)** VCO2 and **(b)** VO2 in DDAH1^Ad-/-^ mice compared to controls. **(c)** Cumulative food intake over the 72 hours. **(d)** Total ambulatory counts (Total XY and Z plane counts). Grey blocks = Dark period. ***P<0.001, Analysis by two-way ANOVA, N=4.

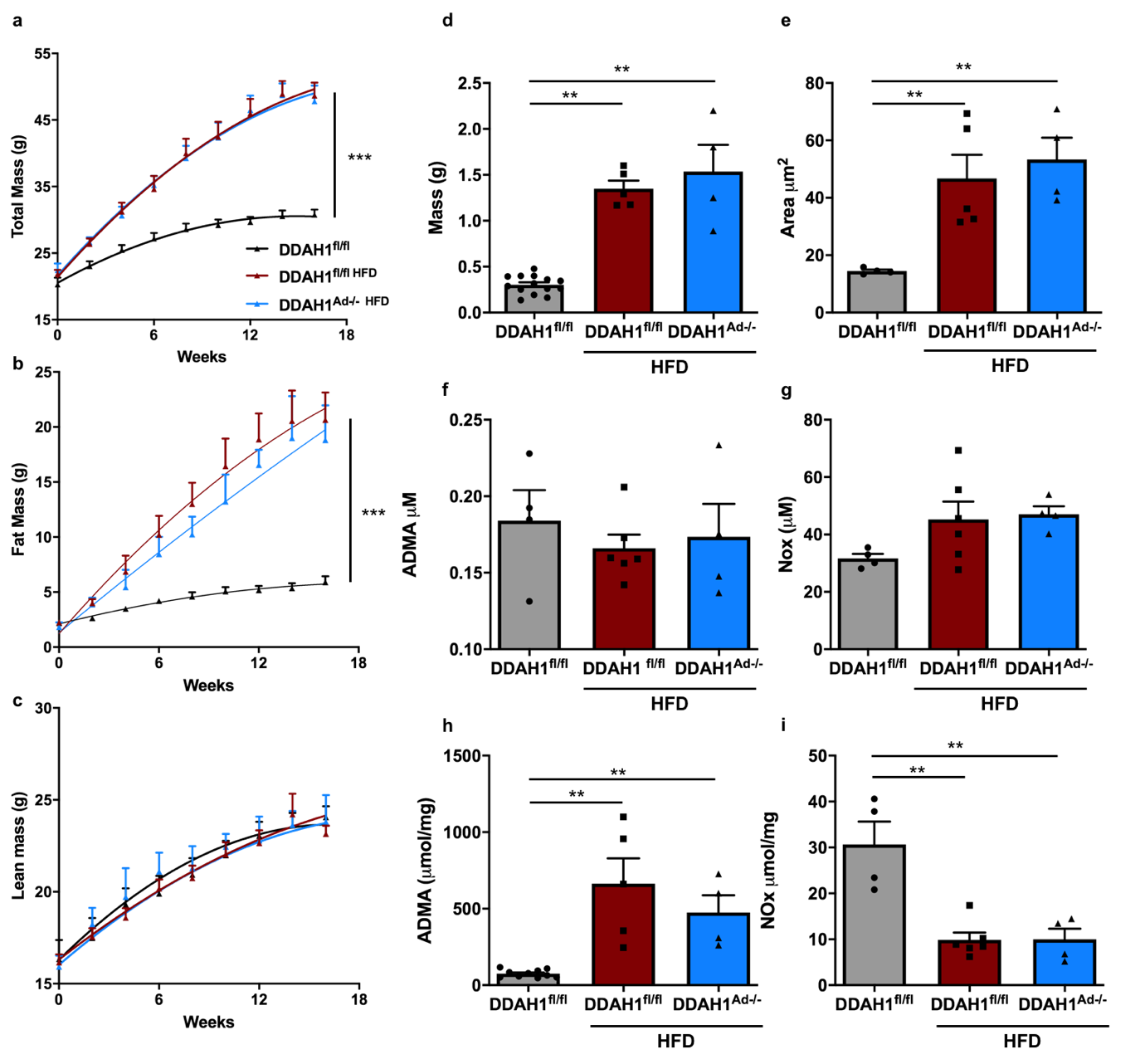

**Supplementary Fig. 5: DDAH1 adipocyte specific deletion does not affect fat mass on a high fat diet; adipocyte ADMA is significantly increased following high fat feeding.** Wt and DDAH1^Adi-/-^ mice were placed on a 70% calories from fat diet from 6 weeks of age for 16 weeks with **(a)** total, **(b)** fat and **(c)** lean mass assessed. Epididymal fat pads wed-ire isolated and assessed for **(d)** mass and **(e)** adipocyte size. ADMA and NOx concentrations in plasma **(f and g)** and isolated adipocytes **(h and i)**. Analysis by two-way ANOVA **(a-c)** and one way ANOVA **(d-i)**; ** p<0.01, ***p<0.001.

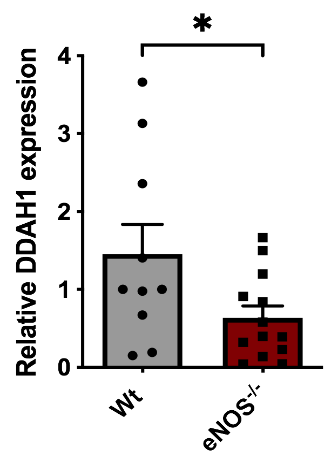

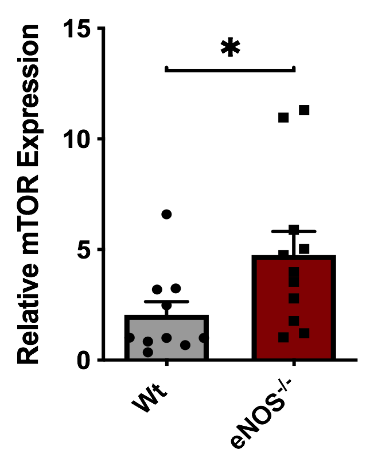

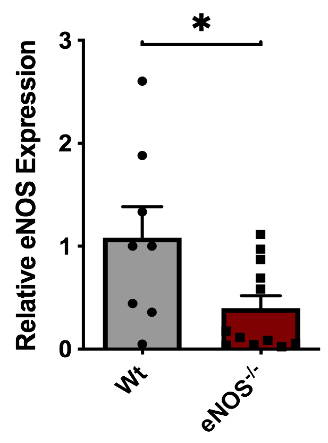

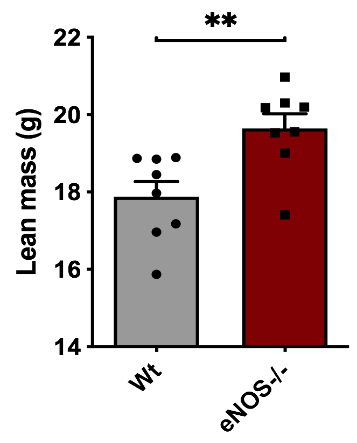

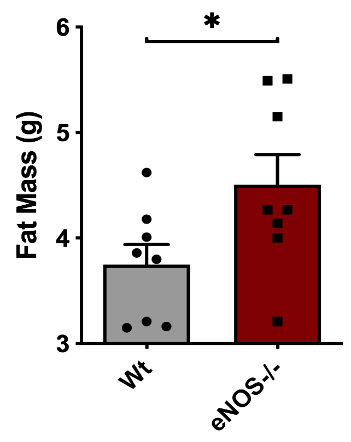

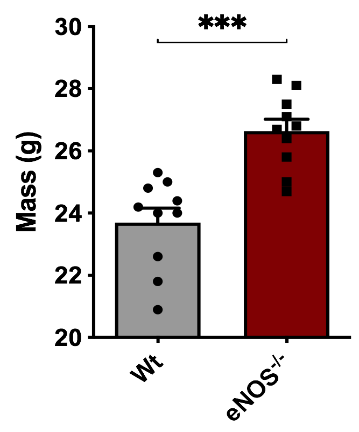

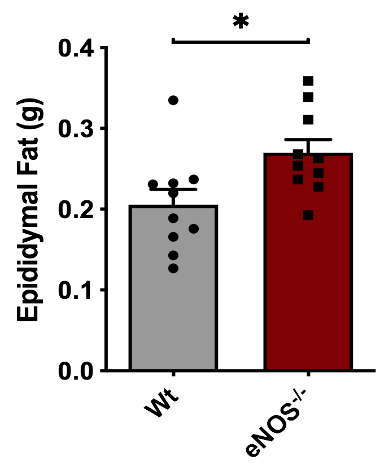

**a**

**c**

**e**

**f**

**d**

**b**

**g**

**Supplementary Fig. 6 : DDAH1 adipocyte expression is downregulated in eNOS^-/-^ mice with enlarged visceral fat.** Male eNOS^-/-^ mice were analysed at 22 weeks of age for **(a)** mass, **(b)** fat mass, **(c)** lean mass and **(d)** isolated epididymal adipose tissue. Following primary adipocyte isolation relative expressions of **(e)** eNOS, **(f)** DDAH1 and **(g)** mTOR were determined by qPCR analysis. Analysed by Mann-Whitney test * p<0.05, ** p<0.01, ***p<0.001

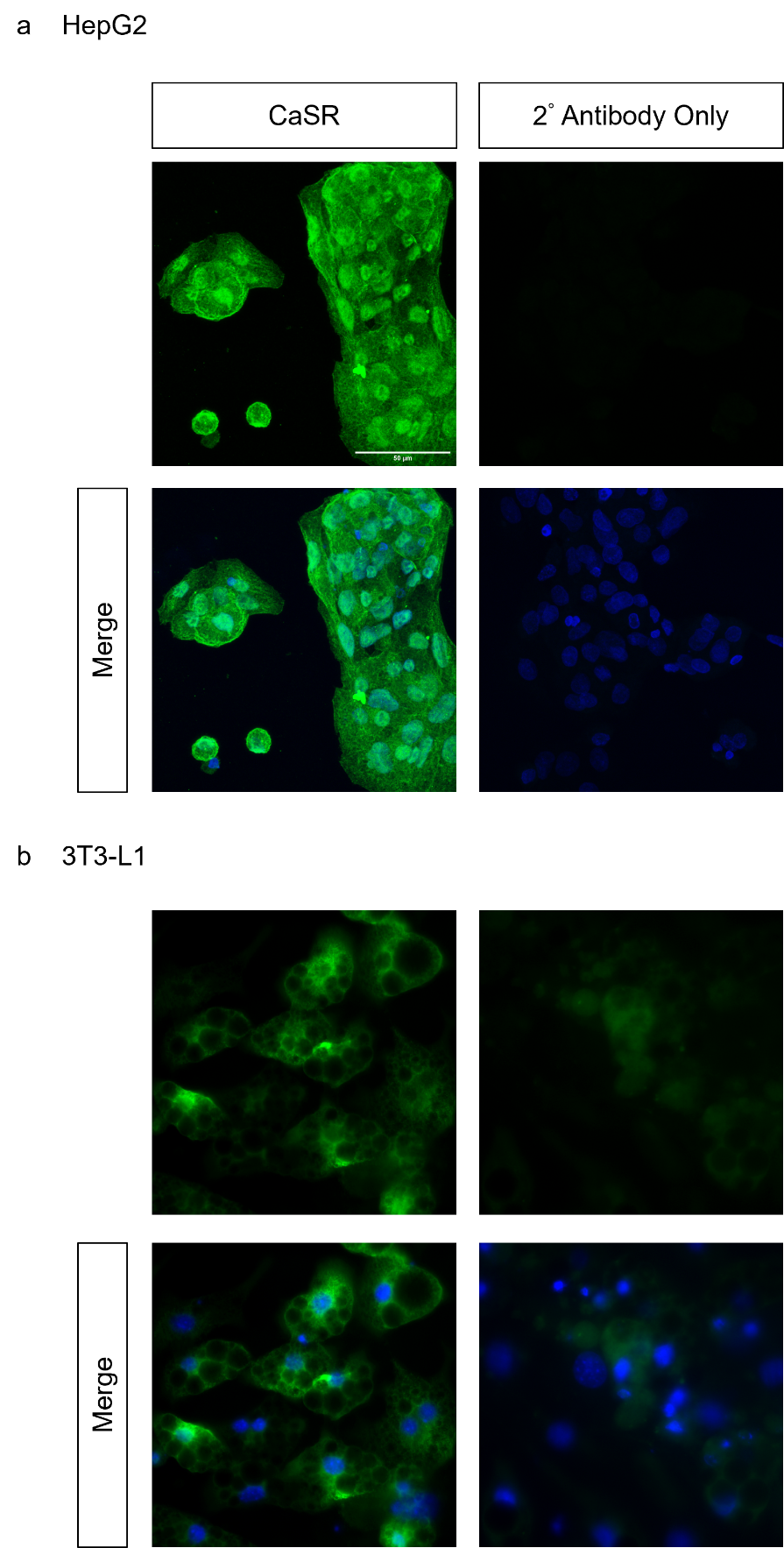

**Supplementary Figure 7: CaSR is expressed in both HepG2 and 3T3-L1 cells.** HepG2 **(a)** and **(b)** 3T3-L1 cells were stained for CaSR (green). DAPI (blue) was used to stain nuclei. Secondary antibody controls were used to confirm specificity of fluorescence.
